# Convergent inactivation of the skin-specific C-C motif chemokine ligand 27 in mammalian evolution

**DOI:** 10.1101/526145

**Authors:** Mónica Lopes-Marques, Luís Q. Alves, Miguel M. Fonseca, Giulia Secci-Petretto, André M. Machado, Raquel Ruivo, L. Filipe C. Castro

## Abstract

The appearance of mammalian-specific skin features was a key evolutionary event contributing for the elaboration of physiological processes such as thermoregulation, adequate hydration, locomotion and inflammation. Skin inflammatory and autoimmune processes engage a population of skin-infiltrating T cells expressing a specific C-C chemokine receptor (CCR10), which interacts with an epidermal CC chemokine, the skin-specific C-C motif chemokine ligand 27 (CCL27). CCL27 is selectively produced in the skin by keratinocytes, particularly upon inflammation, mediating the adhesion and homing of skin-infiltrating T cells. Here, we examined the evolution and coding condition of *Ccl27* in 112 placental mammalian species. Our findings reveal that a number of open reading frame inactivation events such as insertions, deletions, start and stop codon mutations, independently occurred in Cetacea, Pholidota, Sirenia, Chiroptera, and Rodentia, totalizing 18 species. The diverse habitat settings and life-styles of *Ccl27*-eroded lineages probably implied distinct evolutionary triggers rendering this gene unessential. For example, in Cetacea the rapid renewal of skin layers minimizes the need for an elaborate inflammatory mechanism, mirrored by the absence of epidermal scabs. Our findings suggest that the convergent and independent loss of *Ccl27* in mammalian evolution concurred with unique adaptive roads for skin physiology.

## Introduction

The mammalian skin performs a plethora of biological functions, including that of acting as a protective barrier from external harmful insults, such as invading pathogens and noxious stimuli. In this context, the role of the immune system is fundamental, involving a coherent and highly coordinated network of innate and adaptive components to ensure an adequate response to ensure homeostasis [1, 2]. Chemokines, a superfamily of polypeptides, are central in the unfolding of immune and inflammatory responses, serving as chemoattractant signals that drive the movements of immune cells in response to stimuli. In the skin, a tissue-specific T cell homing chemokine has been described. Initially named as the cutaneous T cell-attracting chemokine, C-C motif chemokine ligand 27 (CCL27, also known as ESkine, ALP, ILC or ILRα *locus* chemokine [3-5]) plays a central role in the skin homing process [6]. *Ccl27* maps to human chromosome 9 in a tandem gene arrangement with two other chemokines, *Ccl19* and *Ccl21* and presents two alternative transcripts, yielding secreted and intracellular forms (Figure 1) [3, 7]. The latter, designated PESKY, includes a different exon 1 and acts as an intracellular chemokine (Figure 1) [7]. PESKY transcripts may be found in various mucosal tissues [8, 9], but CCL27 secretion is mostly restricted to skin keratinocytes, having a critical role in skin homeostasis [6, 10]. To provide a skin-specific cue to attract memory T cells in normal or inflamed skin, CCL27 specifically binds to the CCR10 receptor *in vivo* [6, 10]. While CCL27 is exclusive towards CCR10 receptor, other chemokines such as CCL8 also bind CCR10 [6, 9, 10].

**Figure 1:**
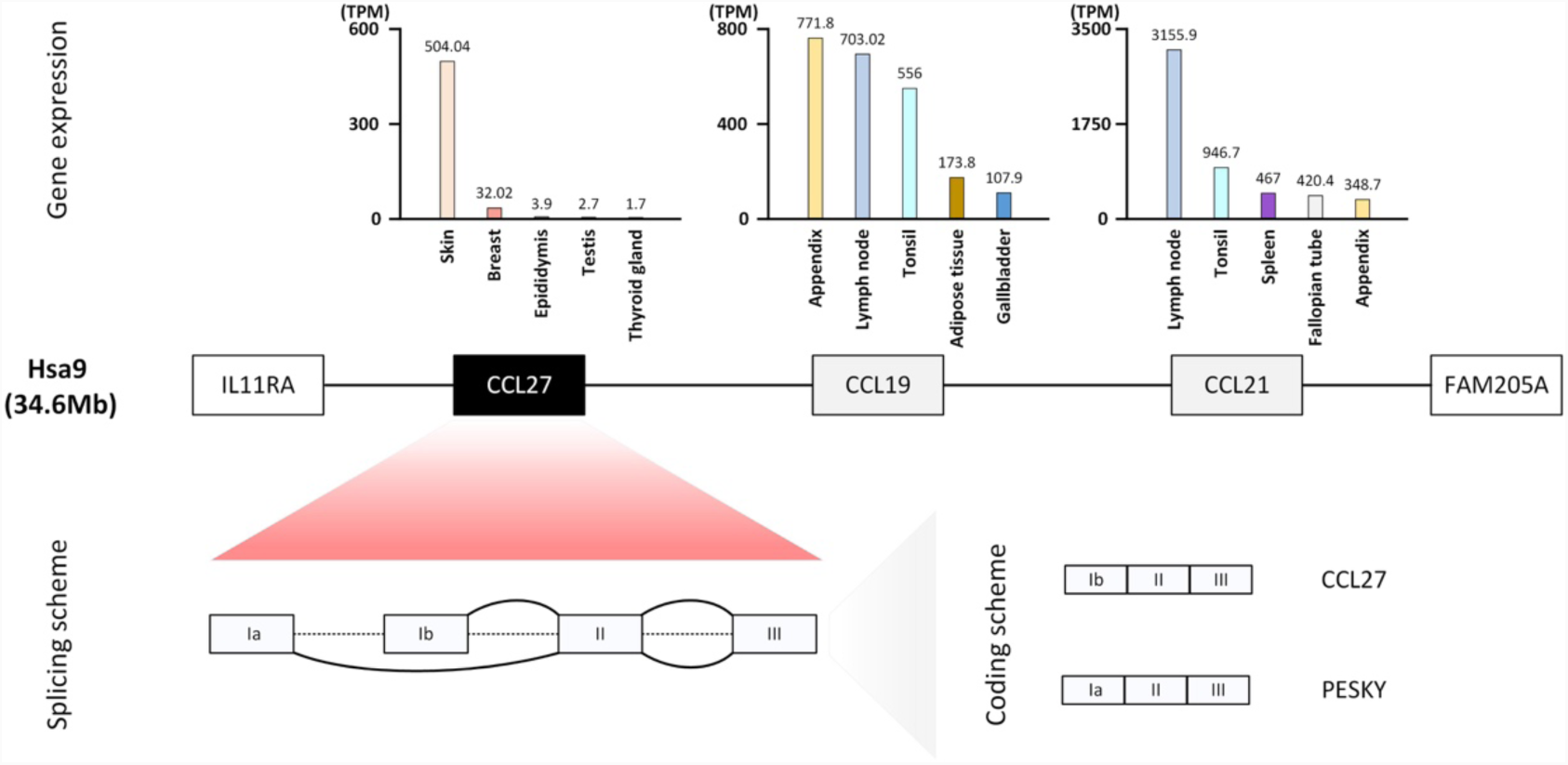
The human orthologue *Ccl27* gene expression, genomic *locus* and structure. In the centre, the genomic region of *Homo sapiens* at chromosome 9, containing *Ccl27* gene (in black box) and tandem gene duplicates, *Ccl19* and *Ccl21* (grey boxes). The corresponding *Ccl* gene expression data was retrieved directly from the Human Protein Atlas (https://www.proteinatlas.org/). Only five tissues with the highest values of transcripts per million (TPM) are represented. Bottom figure represents *Ccl27* gene structure and alternative splicing producing two transcripts: CCL27 and PESKY.

While a number of key morpho-functional skin components have been conserved throughout mammalian evolution, specific lineages experienced secondary episodes of phenotypic simplification or/and elaboration (e.g. [11, 12]). In this context, Cetacea offer an illustrative example, with the exclusive aquatic dependence underscoring unique anatomical signatures (e.g. [12, 13]). For example, their skin is smooth with no pelage, presenting a thick stratum corneum, while the upper layers of the epidermis are not fully cornified [14-16]. Moreover, to improve smoothness and reduce drag, Cetacea skin is rapidly renewed [15, 17, 18]. This intensive cellular replacement and epidermal thickness reduce scab formation and the risk of pathogen invasion [14, 16, 18]. Accordingly, skin inflammation is apparently reduced in Cetacea [18]. The underlying genomic events connected with skin repair mechanisms and whether other mammalian lineages display similar traits is presently unknown. The growing number of full genome sequences currently available have provided valuable insights into the role of gene loss as the foundation for phenotypic alterations and consequently on the perception of adaptive landscapes [11, 12, 19-22]. Given the key role of CCL27 in the process of skin inflammation, we hypothesized that the *Ccl27* coding sequence might be compromised in Cetacea as suggested by the overall skin inflammatory physiology observed in this lineage [15, 18].

## Results and Discussion

To investigate the distribution and annotation tags of the *Ccl27* gene in mammals, we scrutinized a total of 114 selected mammalian genomes available at NCBI and Ensembl genome browsers (supplementary table 1). This search retrieved 14 *Ccl27* annotations tagged as “low-quality” (LQ) and uncovered 9 species with no *Ccl27* gene annotation (supplementary table 1). Next, we investigated the genomic sequences corresponding to the *Ccl27* LQ annotations to determine the CDS through manual annotation. This step revealed coding *Ccl27* genes, tagged as LQ for the following species: *Saimiri boliviensis* (black-capped squirrel monkey), *Galeopterus variegatus* (Sunda flying lemur), *Peromyscus maniculatus bairdii* (North American deer mouse), *Loxodonta Africana* (African bush elephant), and *Chrysochloris asiatica* (Cape golden mole). Also, the analysis of the genomic sequence corresponding to the *Ccl27 locus* in *Ochotona princeps* (American pika) showed that the missing annotation in this species is most probably due to poor genome coverage in this *locus* (not shown). Importantly, all cetacean species analysed presented sequences tagged as LQ or no *Ccl27* annotation. This impelled us to further explore other cetacean species with unannotated genomes: *Balaenoptera bonaerensis* (Antarctic minke whale), *Eschrichtius robustus* (gray whale), *Balaena mysticetus* (bowhead whale) and *Sousa chinensis* (Indo-Pacific humpback dolphin).

### *Ccl27* gene sequence contains inactivating mutations in Cetacea

Annotation of collected cetacean genomic sequences revealed *Ccl27* gene erosion across all analysed species (Figure 2A). In detail, gene sequence examination in Odontoceti showed a non-disruptive insertion of a codon in exon 2 in all species with the exception of *Lipotes vexillifer* (Yangtze river dolphin). In addition, in exon 2, a frameshift mutation (deletion of 1 nucleotide) was identified and validated by sequence read archive (SRA) analysis in *Physeter catodon* (sperm whale, supplementary material 1). A conserved premature stop codon was found in *Orcinus orca* (orca) and *Lagenorhynchus obliquidens* (Pacific white-sided dolphin), as well as a non-conserved premature stop codon in *L. vexillifer.* These observations were confirmed in *O. orca* and *L. obliquidens* through SRA analysis (supplementary material 1). In *L. vexillifer* exon 2 also presented a frameshift mutation (4 nucleotide deletion) and the loss of the canonical splice site (GT>CC). Next, in exon 3 a conserved premature stop codon was identified in all Odontoceti (Figure 2B, supplementary material 1). A frameshift mutation by deletion was identified in all species apart from *P. catodon*, which in turn shows a frameshift mutation before the identified stop codon (Figure 2B).

**Figure 2:**
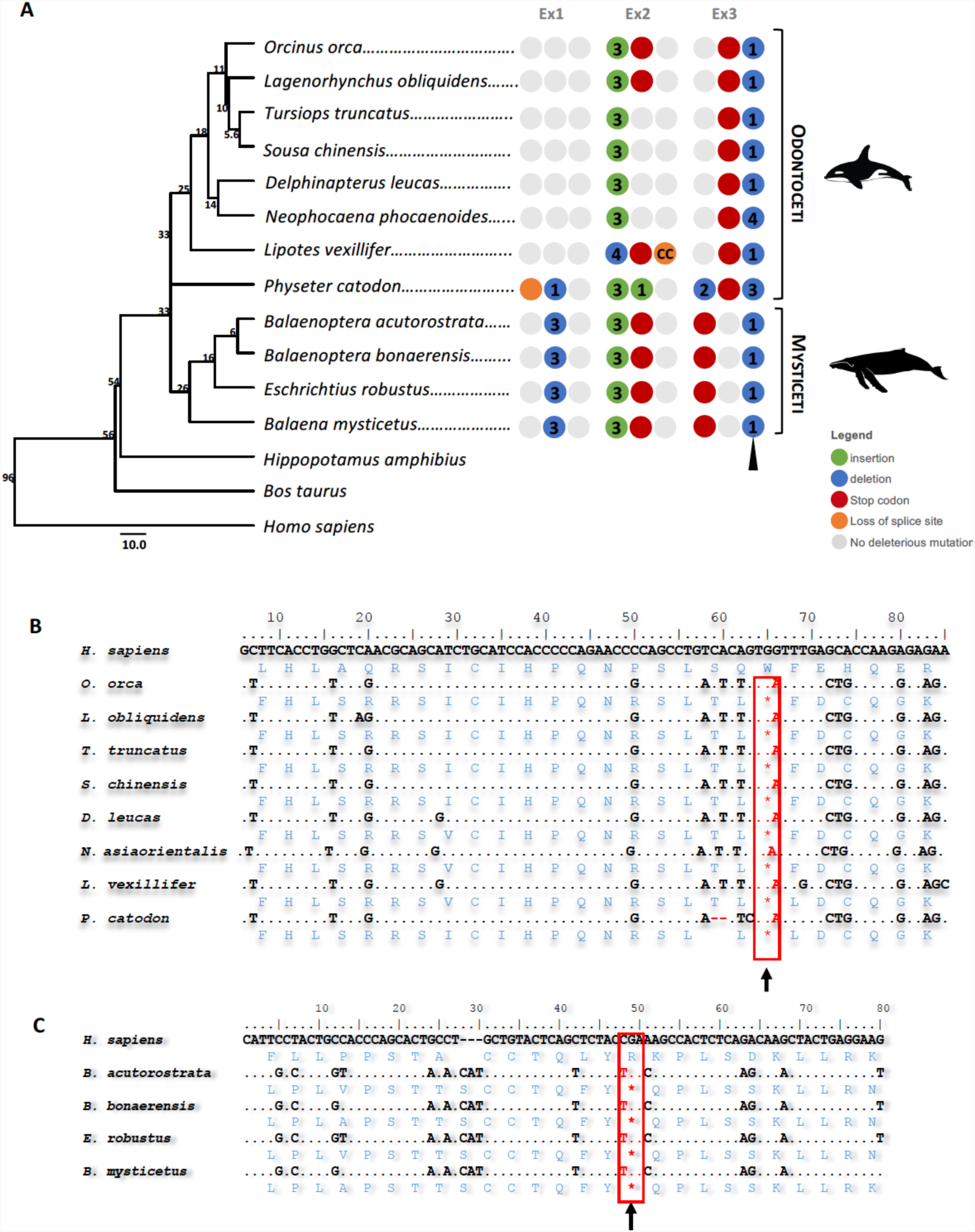
**A**- Schematic representation of the *Ccl27* gene and identified mutations in Cetacea, each group of 3 circles represents one exon, red represents stop codon; orange, non AG-GT splice site; blue deletion and green nucleotide insertion; numbers at tree nodes indicate million years. Number in the circles indicate number of nucleotides inserted or deleted and dark grey circles represent regions or exon not found. **B**- Sequence alignment of the identified premature stop codon in exon 3 of Odontoceti. **C**- Sequence alignment of the identified premature stop codon in exon 2 of Mysticeti.

Regarding the Mysticeti, all identified mutations were conserved across all 4 analyzed species (Figure 2A and 2C). Non-disruptive mutations consisting in the deletion and insertion of 1 codon were identified in exon 1 and exon 2, respectively. Also, two conserved premature stop codons were identified in exon 2 and exon 3 (the former was validated by SRA, supplementary material 1), which were followed by a 1 nucleotide deletion identified in all analyzed species (Figure 2C black arrow). Interestingly, this 1 nucleotide deletion is conserved among all cetacean species (supplementary material 2), suggesting that *Ccl27* pseudogenization preceded the divergence of Odonticeti and Mysticeti.

### Transcriptomic analysis supports *Ccl27* gene erosion in Cetacea

To further scrutinize the functional condition of *Ccl27*, we next analyzed multi-tissue RNA-Seq projects available at NCBI for 6 cetacean species: *Tursiops truncatus* (common bottlenose dolphin), *Delphinapterus leucas* (beluga whale), *Neophocaena asiaeorientalis* (finless porpoise), *P. catodon* (sperm whale), *Balaenoptera acutorostrata* (common minke whale), and *B. mysticetus* (supplementary table 2).

Overall, RNA-Seq analysis revealed a considerably low number of *Ccl27* mRNA reads across all the 6 species, especially in *N. asiaeorientalis* (Figure 3). Moreover, for the remaining species we observed a substantially high proportion of reads spanning adjacent exonic and intronic regions, exon-intron reads, versus spliced reads, connecting contiguous exons and containing no intronic remnants, especially in the case of *T. truncatus* (121 exon-intron reads against 3 spliced reads). In the later, the higher number of skin-specific sequencing runs available for this species, compared to the remaining ones (25 skin-specific sequencing runs in *T. truncatus vs.* an average of 6.6 skin-specific sequencing runs per species), probably explains the variation in the number of exon-intron reads (see supplementary table 2). As we observed a specific case with a considerably distinct ratio of exon-intron reads/spliced reads amongst the remaining species, namely *B. mysticetus* (49 spliced reads vs 52 exon-intron reads), we decided to further verify the presence of ORF disruptive mutations in the produced transcripts of *Ccl27* in each of the referred species. We were able to detect at least one premature stop codon in the transcripts of the analysed 6 cetacean species (see supplementary material 3), revealing that *Ccl27* transcripts contained the genome predicted ORF mutations (Figure 3).

**Figure 3:**
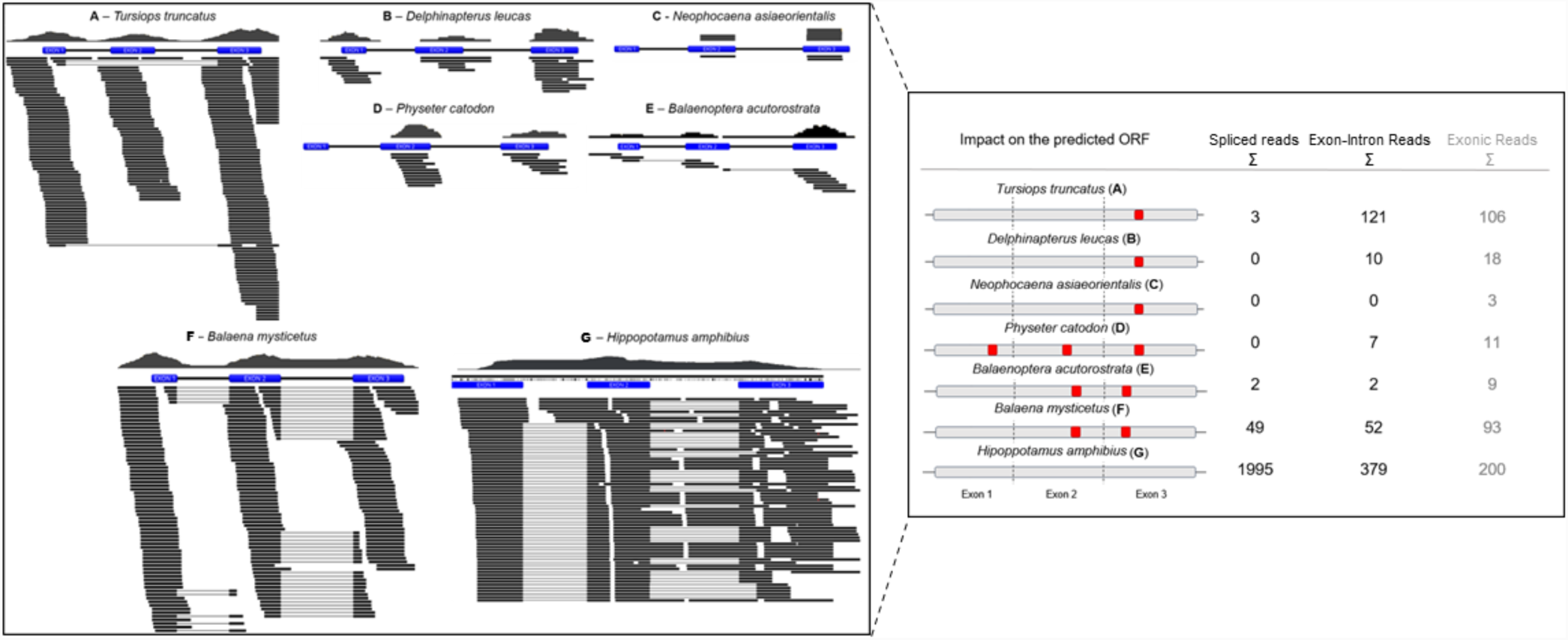
Gene expression of *Ccl27* across Cetacea species. In the left box: mapping of the NCBI Sequence Read Archive (SRA) recovered multi-tissue RNA-Seq reads (black) for each of the 7 represented species against the corresponding *Ccl27* annotated gene (blue). Right box: impact of the annotated mutations in the open reading frame (ORF) of the *Ccl*27 gene. Premature stop codons are represented with a red squared marker at the corresponding exon. Overall count of RNA-Seq mapped reads for each specie. Reads are classified into spliced reads (reads spanning over two different exons), exon-intron reads (reads containing exonic and intronic sequence) and exonic reads (reads fully overlapping exonic regions).

The conserved mutational pattern observed between Odontoceti and Mysticeti suggests that *Ccl27* inactivation occurred in the Cetacea ancestor. To further survey and estimate the approximate timing of *Ccl27* loss in Cetacea, we next investigated the genome and the skin transcriptome of the extant sister clade of the Cetacea, the Hippopotamidae. The current version of the *H. amphibius* genome, available at NCBI, is fragmented and unannotated (GCA_002995585.1). However, we were able to deduce the full coding ORF of the *Ccl27* gene orthologue in *H. amphibius*, and without any intervening inactivating mutation (Figure 3). Furthermore, by examining a skin-specific transcriptome we identified a very high proportion of spliced/exon-intron mRNA reads (1995 spliced reads against 379 exon-intron reads), a clear indication that the gene is functional in this species (Figure 3).

### *Ccl27* is eroded in other non-cetacean mammals

We next investigated the uniqueness of *Ccl27* inactivation in other mammalian lineages with absent or LQ annotations. Our initial analysis revealed that several genes annotated as LQ were in fact coding. For example, the analysis of the retrieved genomic sequence for *L. africana* revealed poor genome coverage in the *Ccl27 locus*. However, blast search of the whole genome sequence recovered a genomic scaffold (NW_003573426.1:65683000-65687000) that contained an intact *Ccl27* gene sequence. Yet, in the case of LQ tagged *Ccl27* from *Hipposideros armiger* (great groundleaf bat), *Trichechus manatus* (West Indian manatee), *Heterocephalus glaber* (naked mole rat), and *Manis javanica* (Sunda pangolin) the analysis and manual annotation of the corresponding genomic sequence revealed a number of ORF disrupting mutations (Figure 4A). Our findings were further supported by searching the available unannotated genomes of *Manis pentadactyla* (Chinese pangolin) and *Rhinolophus sinicus* (Chinese rufous horseshoe bat), which after annotation also presented a non-coding *Ccl27* ORF (Figure 4A). Briefly, *Ccl27* annotation in *H. armiger* revealed a premature stop codon in exon 3 followed by 1 nucleotide insertion, and in *R. sinicus* a single premature stop codon was identified in exon 2 (all confirmed by SRA search Supplementary material 4). In the Pholidota *M. javanica* and *M. pentadactyla* a shared frameshift mutation in exon 2 was identified (validated by SRA in *M. javanica* Supplementary material 4). Additionally, *M. pentadactyla* presents a premature stop codon in exon 2 while *M. javanica* presents a premature stop codon in exon 3 preceded by a 1 nucleotide frameshift mutation. In rodentia, the *Ccl27*gene annotation in *H. glaber* revealed a missing start codon in exon 1 combined with a premature stop codon in exon2, and finally in *T. manatus Ccl27* gene annotation uncovered a premature stop codon in exon 3 (stop codons validated by SRA Supplementary material 4).

**Figure 4:**
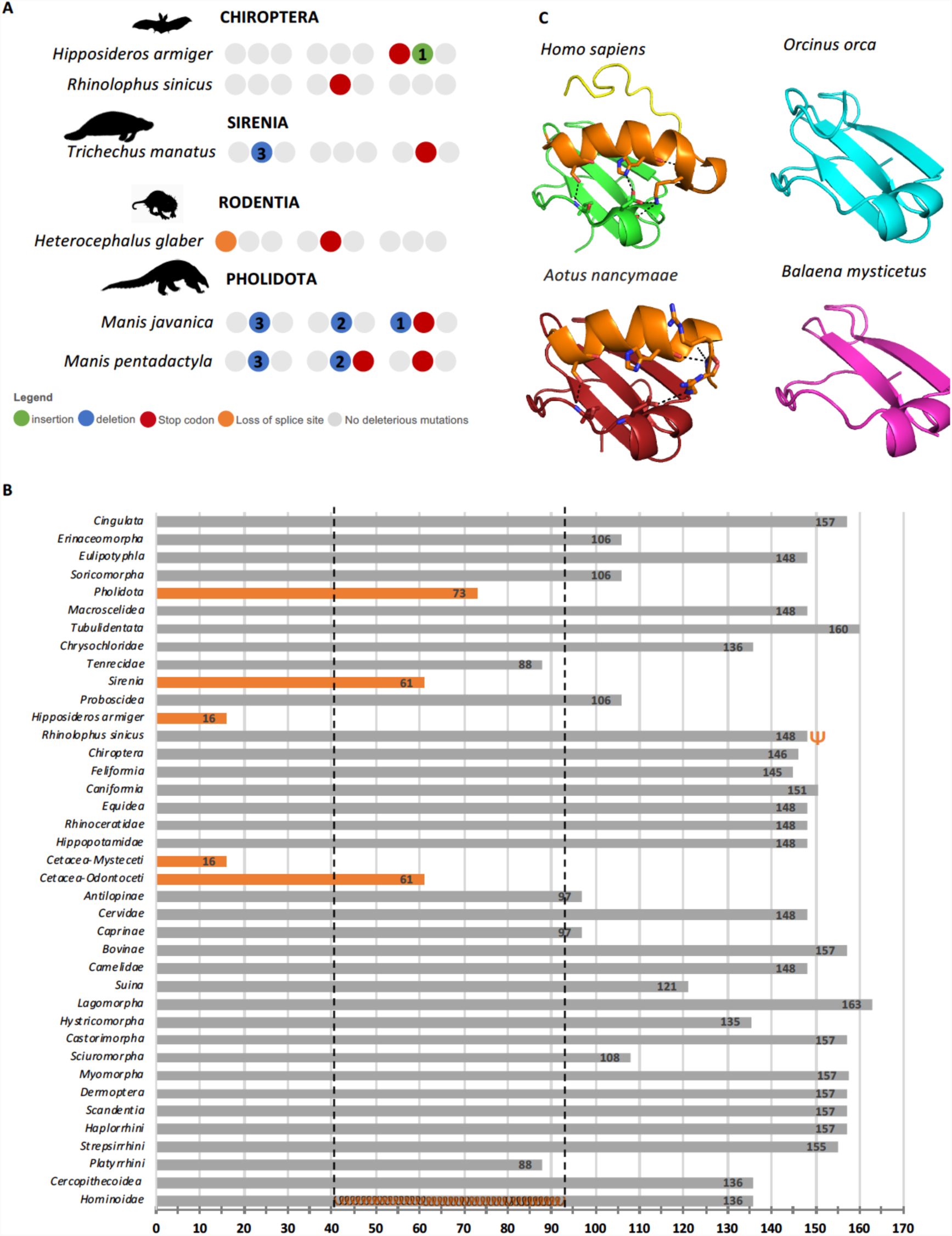
**A**- Gene annotation of *Ccl7* in non-cetacean mammals. **B-** Analysis of exon3 length in nucleotides, orange bars highlights species with severe exon 3 truncation, orange helix in Hominoidae bar corresponds to extension C-terminal α-helix in human crystal structure (2KUM). **C-** Comparative analysis of the human crystal structure 2KUM (green) and calculated homology models in red *Aotus nancymaae*, blue *Orcinus orca* and magenta *Balaena mysticetus*. Structural features highlighted in human in orange terminal α-helix, in yellow disordered terminal region.

### Exon 3 length reduction parallels gene inactivation

The survey of 112 placental mammalian *Ccl27* CDS exposed a variable C-terminal length in different mammalian species. Thus, we compared the predicted length of exon 3 in the annotated pseudogenes regardless of prior ORF disrupting mutations (Figure 4B). This analysis showed that all annotated pseudogenes were severely truncated in exon 3 with the exception of *R. sinicus.* Also, the analysis of the observed truncations in the overall structure of CCL27 using homology modelling for *O. orca* and *B. mysticetus* showed that premature stop codons occur early in the C-terminal α-helix. CC chemokines present a highly conserved quaternary structure characterized by disordered N-terminal region followed by a 3_10_-helix, 3 antiparallel β-strands, followed a C-terminal α-helix and ending with a disordered stretch of positive residues [23]. Interestingly, the C-terminal region, specifically the disordered region, is a feature that differentiates CCL27 from the majority of CC chemokines, and has been shown to be involved in nuclear import [9]. In agreement, both *Ccl27* transcript variants, the intracellular chemokine PESKY and the internalized CCR10-bound CCL27, target the cell nucleus, modulating morphology and motility via transcriptional modification [4, 8]. Moreover, the remaining mammals including Tenricidae, Antilopinae, Caprinae, Platyrrhini, exhibit a sequence deletion pattern at the end or shortly after the α-helix (Figure 4B), which implies the loss of the final C-terminal disordered region involved in nuclear targeting. Yet, contrarily to the annotated pseudogenes, the coding *Ccl27 Aotus nancymaae,* which presents the shortest exon 3, still conserves the full α-helix which has been reported to stabilize the overall fold [23] (Figure 4C). The biological significance of this plasticity remains to be studied.

### *Ccl27* gene loss correlates with alternative protection and healing programs

Together, our analysis indicates that *Ccl27* is most likely non-functional in all of the examined cetacean species. Inactivating mutations are also present in species of Pholidota, Sirenia, Chiroptera, and Rodentia. Even if a full phenotypic description of mouse knock-out (KO) for this gene is presently unavailable, the initial data suggests a decrease of the T cell population in intact skin [24]. On the other hand, constitutive production of keratinocyte CCL27 enhanced the inflammatory response in mice [25]. In agreement, chronic inflammatory skin diseases, such as atopic dermatitis and psoriasis, are characterized by increased serum levels of the T cell attracting chemokine CCL27 [10], while CCL27*-*neutralizing antibody treatment reduced skin inflammation in a transgenic animal model [26]. Thus, upon insult or infection, *Ccl27* KO would likely show an attenuated inflammatory response in the skin. This hypothesis remains to be verified.

Nevertheless, it could be argued that the premature stop codon in exon 3 of *Ccl27* in *T. truncatus, D. leucas, S. chinensis* and *N. asiaorientalis* could still encode a functional shorter isoform. Yet, RNA-Seq transcriptome and structural analysis supports a different interpretation. Since *Ccl27* prime expression site is the skin, we analysed the available skin and multi-tissue RNA-transcriptomes from Cetacea and found two distinct scenarios. First, in the majority of the species, RNA-Seq searches recovered reads covering exon-intron. Second, in *B. mysticetus* the recovered RNA-Seq reads presented a higher number of spliced reads. However, in both cases, detailed analysis of the collected reads confirmed the presence of the previously identified ORF-disrupting mutations. Thus, we suggest that these mRNA mature sequences do not translate into a functional protein. In addition, previous studies, addressing the loss of visual opsins in Chiroptera, highlighted possible discrepancies between gene integrity and protein production: further suggesting post-transcriptional mechanisms as regulators of evolutionary gene silencing [22].

Our analysis strongly supports that *Ccl27* gene pseudogenization compromises both canonical CCL27 and PESKY transcripts. This is in accordance with previous findings reporting a distinct inflammatory and wound healing program in cetacean skin [18, 27]. Interestingly, scar-less and low inflammation wound repair has also been reported in several mammalian foetus, including human, as well as in human adult oral mucosa [28, 29]. Both observations might correlate with decreased or null CCL27 secretion. In fact, embryonic keratinocytes are more proliferative and less immunogenic than adult cells, inhibiting T cell proliferation [30]. Similarly, oral mucosa exhibits rapid wound healing due to accelerated re-epithelialization [28]. In wounded oral mucosa, the overall expression of *Ccl27* is also downregulated when compared to wounded skin [28]. Thus, in scab-less and low inflammation wound repair, increased epithelial renewal seems to parallel the downregulation or absence of CCL27 secretion. Yet, in oral mucosa the possible maintenance of PESKY could participate in the healing circuitry by stimulating cell migration and proliferation.

Additionally, we found convergent inactivation of *Ccl27* in other non-cetacean mammalian species: namely in pangolins (*M. javanica* and *M. pentadactyla*), in the naked mole rat (*H. glaber*), in the sirenian *T. manatus,* and in two Chiroptera (*H. armiger* and *R. sinicus*). Curiously, with the exception of Chiroptera, these species share some of the distinctive features of Cetacea skin: for example, the hairless phenotype is observed in Pholidota, Sirenia, and naked mole rat, increased epidermal thickness is observed in Sirenia and naked mole rat and Sirenia skin is also smooth [14, 31]. The diversity of skin phenotypes along with the scarce information regarding species-specific inflammatory and wound healing programs, hampers the anticipation of the possible outcomes of *Ccl27* pseudogenization. Nonetheless, the available information suggests that *Ccl27* erosion occurred in species exhibiting singular epidermal renewal, or even protective mechanisms or structures, reducing the need for CCL27-dependent inflammatory processes. For instance, pangolins present a protective armour with keratin-derived scales, which was suggested to reduce epithelial immune requirements [32, 33]. In agreement, pseudogenization of Interferon Epsilon, which confers protection against viral and bacterial infections, was also reported in these species [33]. On the other hand, the naked mole rat abundantly produces high molecular weight hyaluronic acid, suggested to underscore their peculiar longevity and cancer resistance but also contributing to cell motility, rapid wound healing and immunity [31, 34]. Regarding Chiroptera, although their skin is generally similar to most mammalian species, interdigital skin membranes are thinner, and thus more susceptible to damage; yet, interdigital membranes have an enhanced healing capacity [35]. Nonetheless, the inflammatory circuitry of this healing process is still poorly studied. Also, *Ccl27* pseudogenization was only detected in two Chiroptera species. Again, post-translational events could promote CCL27 loss in additional species [22]. In conclusion, our findings reinforce gene loss mechanisms as evolutionary drivers of skin phenotypes in mammals, and correlate *Ccl27* loss with species-specific scar-less and/or low inflammation wound repair.

## Material and Methods

### Sequence retrieval

*Ccl27* coding nucleotide sequences were searched and collected from NCBI for a set of mammalian species representative of the major mammalian lineages (see Supplementary table 1). Searches were performed through tblastn and blastn queries using the human *Ccl27* sequence as reference. Full coding sequences and corresponding genomic sequences were collected, for phylogenetic analysis and gene annotation respectively. Coding sequences were next uploaded into Geneious R7.1.9 curated by removing 5′ and 3′ UTR (untranslated regions) and aligned using the translation align option. Sequence alignment was inspected and exported for phylogenetic analysis. Maximum likelihood Phylogenetic analysis was performed in PhyML3.0 server [36], with best sequence evolutionary model determined using smart model selection [37], and branch support with the aBayes algorithm [38]. The resulting phylogenetic tree was then visualized and analysed in Figtree (Supplementary material 5).

### Gene annotation

For gene annotation the genomic sequence of *Ccl27* annotations tagged as LQ was collected from NCBI. For species with no *Ccl27* annotation (*B. acutorostrata, O. orca* and *R. sinicus*), the genomic sequence ranging from the upstream to the downstream flanking genes was collected. Finally, for species with no annotated genome (*B. bonaerensis, E. robustus, B. mysticetus, H. amphibius* and *M. pentadactyla)*, genomic sequences were recovered through tblastn searches in the whole genome assembly and scaffold corresponding to the highest identity hits were taken. Collected genomic sequences were next loaded to Geneious R7.1.9 for manual annotation as previously described [11, 39]. Briefly, using as reference human and *Bos taurus Ccl27* CDS sequence as reference each individualized exon was mapped on the corresponding genomic sequences using the built-in map to reference tool in Geneious R7.1.9. Aligned regions were manually inspected to verify coding status and identify ORF disrupting mutations (frameshifts, premature stop codon, loss of canonical splice sites). The identified mutations were next validated in at least two independent SRA projects (when available) (see supplementary material 1).

### Transcriptomic Analysis

RNA-Seq analysis was performed to assess the functional condition of *Ccl27* in 6 cetacean species and *H. amphibius*. For each of the 6 cetaceans, using the discontiguous megablast task from Blastn, the *B. taurus Ccl27* coding sequence (CDS) was used as query sequence to recover reads from the totality of the available transcriptomic sequence read archive (SRA) projects available at NCBI. The supplementary table 2 provides an in-depth description of the explored NCBI SRA projects per species. In the case of *H. amphibius*, through megablast from Blastn, the CDS of the annotated gene in the same species was used as query sequence and reads were recovered from the available *H. amphibius* skin transcriptome (accession number PRJNA507170). The collected mRNA reads were mapped against the corresponding annotated gene using the map to reference tool from Geneious R7.1.9. The aligned regions were manually curated, and poorly aligning reads manually removed. Next reads were then classified as spliced reads (reads spanning over two different exons) and exon-intron reads (reads containing intronic sequence). Reads fully overlapping a single exon, exonic reads, were considered inconclusive for this analysis, given that it is infeasible to infer the nature of the corresponding transcript (spliced or unspliced).

### Comparative homology modelling

Comparative homology modelling was performed for *O. orca* representative of Odontoceti, *B. mysticetus* representative of Mysticeti and for *Aotus nancymaae* (Platyrhini) representing a coding CDS with short C-terminal. Predicted CDS sequences of *O. orca* and *B. mysticetus* were determined using the annotated exons and premature stop codons identified in exon 2 were reverted to the residue observed in *B. taurus*, while mutations in exon 3 were left as observed. Corresponding protein sequences were then next submitted to the SWISS-MODEL [40, 41] for homology modelling using the human CCL27 crystal structure as reference (2KUM) [23]. Resulting models were downloaded and analysed in PyMOL V1.74 [42].

## Acknowledgments

This work was supported by Project No. 031342 co-financed by COMPETE 2020, Portugal 2020 and the European Union through the ERDF, and by FCT through national funds.

## Supplementary Information Legends

**Supplementary Table 1:** Accession numbers of the analysed sequences * tagged low-quality, ^a^ assembled genomes without annotation.

**Supplementary Table 2**: In-depth description of the available transcriptomic NCBI sequence read archive (SRA) projects, scrutinized in the transcriptomic analysis of the 6 represented cetaceans.

**Supplementary Material 1**: SRA validation of the identified mutations in Cetacea

**Supplementary Material 2**: Sequence alignment of *Ccl27* exon 3 from Cetacea, *H. amphibius* and *H. sapiens*.

**Supplementary Material 3**: SRA validation of inactivating mutations of *Ccl27* transcripts in Cetacea.

**Supplementary Material 4**: SRA Validation of identified mutations in other mammals.

**Supplementary Material 5:** Distribution and phylogenetic analysis of coding *Ccl27* in mammals.

