## Supplementary Material 4 for "Convergent inactivation of the skin-specific C-C motif chemokine ligand 27 in mammalian evolution"

### Slide 1
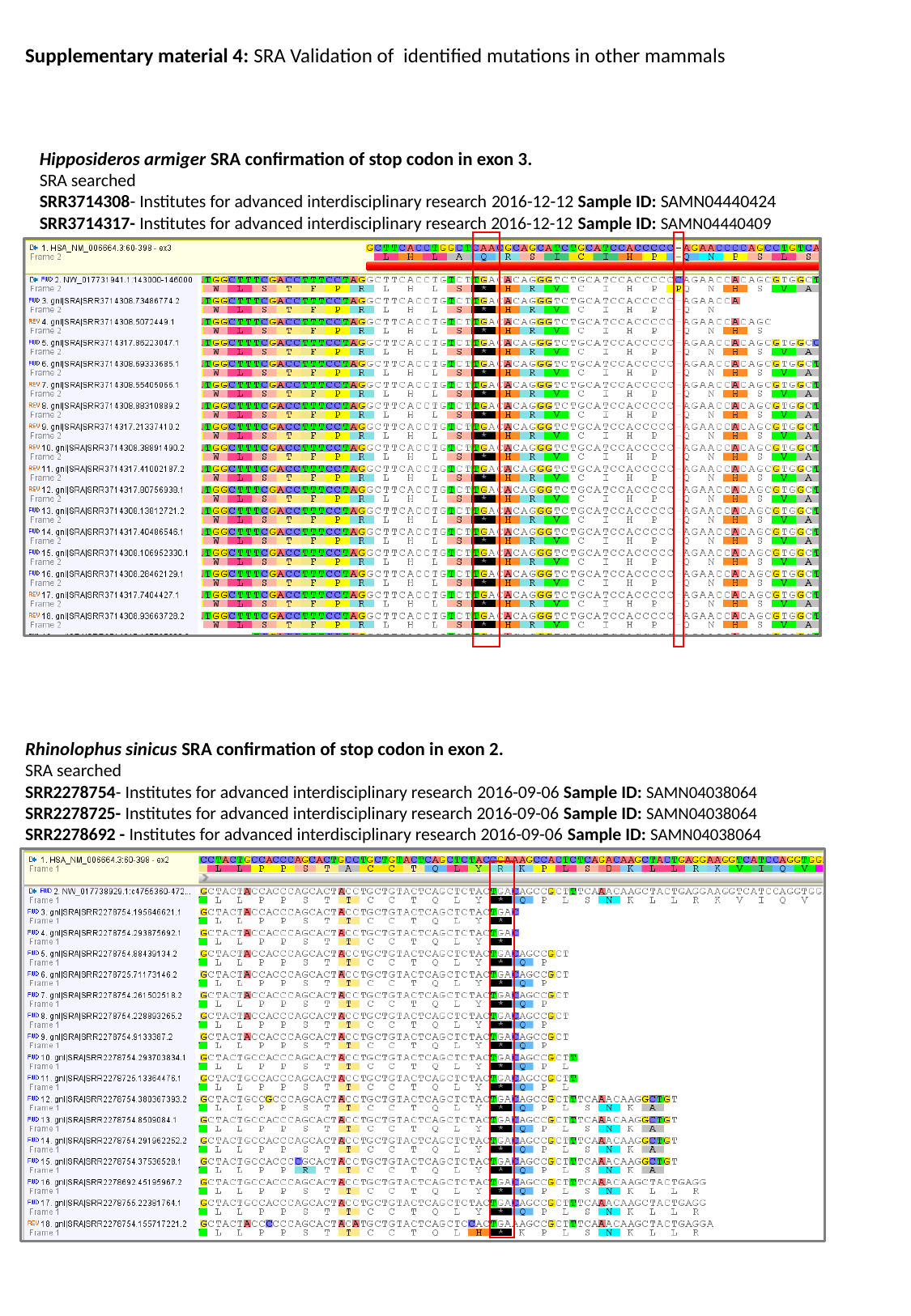

Supplementary material 4: SRA Validation of identified mutations in other mammals
Hipposideros armiger SRA confirmation of stop codon in exon 3.
SRA searched
SRR3714308- Institutes for advanced interdisciplinary research 2016-12-12 Sample ID: SAMN04440424
SRR3714317- Institutes for advanced interdisciplinary research 2016-12-12 Sample ID: SAMN04440409
Rhinolophus sinicus SRA confirmation of stop codon in exon 2.
SRA searched
SRR2278754- Institutes for advanced interdisciplinary research 2016-09-06 Sample ID: SAMN04038064
SRR2278725- Institutes for advanced interdisciplinary research 2016-09-06 Sample ID: SAMN04038064
SRR2278692 - Institutes for advanced interdisciplinary research 2016-09-06 Sample ID: SAMN04038064

### Slide 2
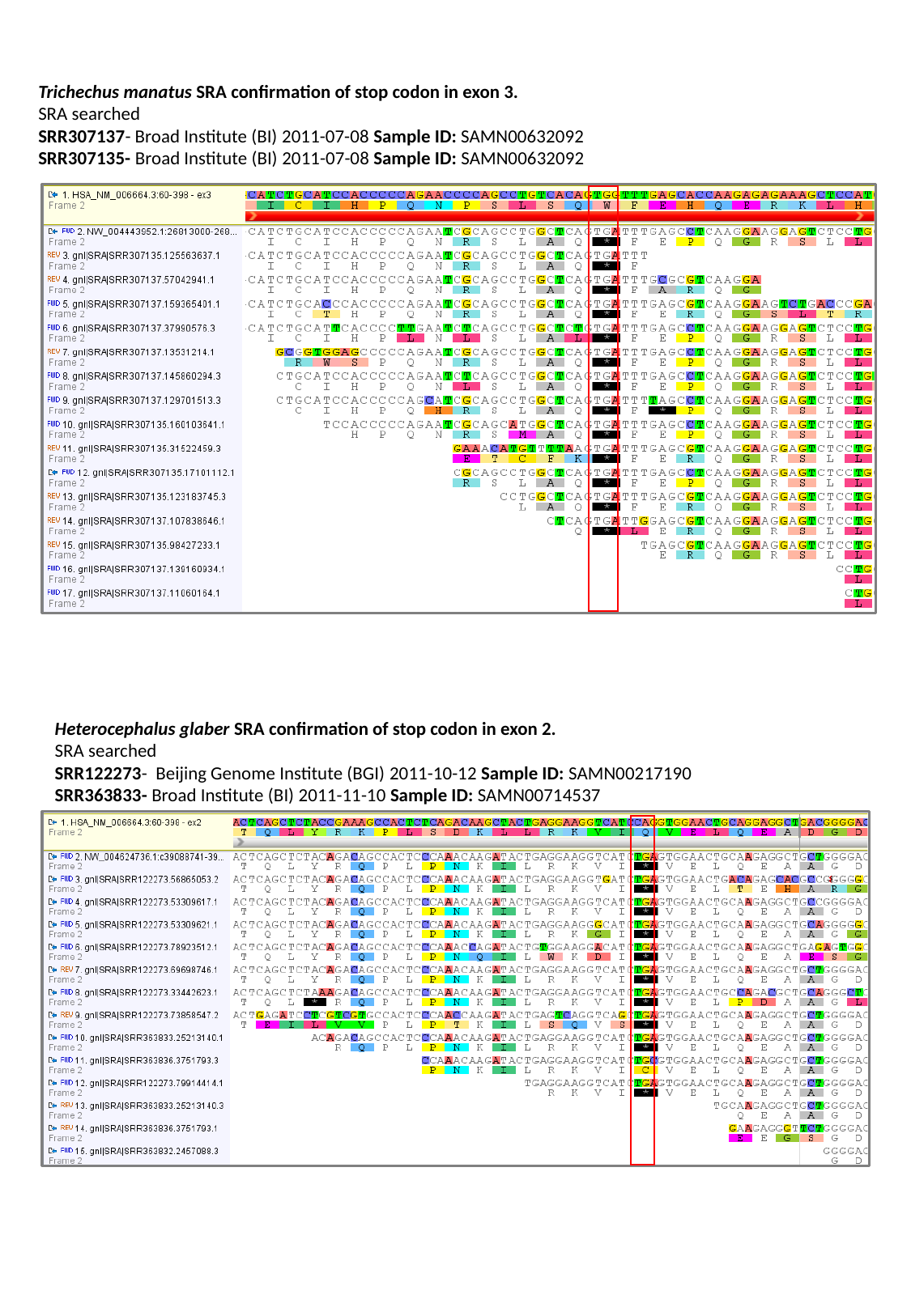

Trichechus manatus SRA confirmation of stop codon in exon 3.
SRA searched
SRR307137- Broad Institute (BI) 2011-07-08 Sample ID: SAMN00632092
SRR307135- Broad Institute (BI) 2011-07-08 Sample ID: SAMN00632092
Heterocephalus glaber SRA confirmation of stop codon in exon 2.
SRA searched
SRR122273- Beijing Genome Institute (BGI) 2011-10-12 Sample ID: SAMN00217190
SRR363833- Broad Institute (BI) 2011-11-10 Sample ID: SAMN00714537

### Slide 3
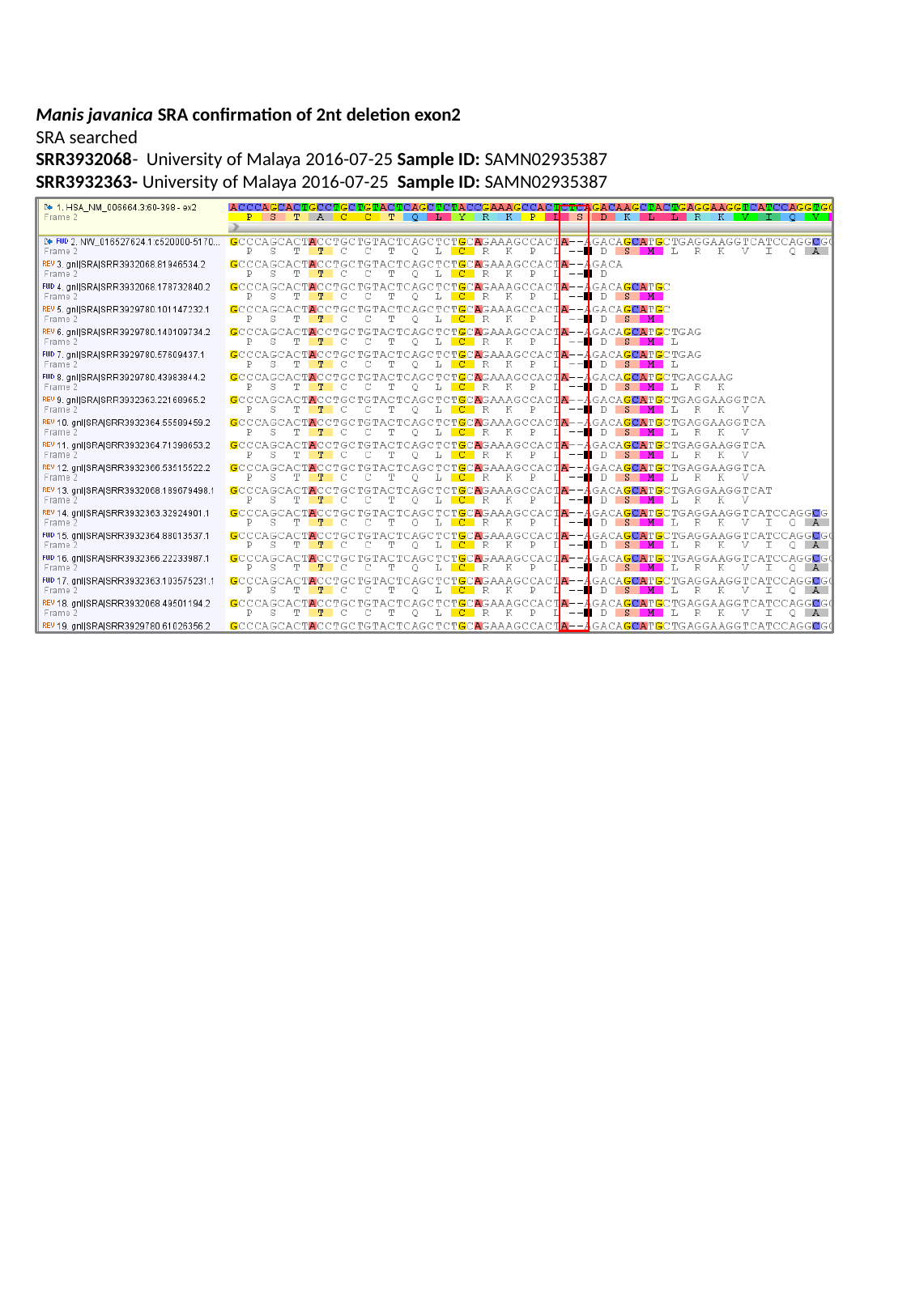

Manis javanica SRA confirmation of 2nt deletion exon2
SRA searched
SRR3932068- University of Malaya 2016-07-25 Sample ID: SAMN02935387
SRR3932363- University of Malaya 2016-07-25 Sample ID: SAMN02935387
