## Supplementary Material 2 for "Convergent inactivation of the skin-specific C-C motif chemokine ligand 27 in mammalian evolution"

**Supplementary material 2**: Sequence alignment of *Ccl27* exon 3 from Cetacea, *H. amphibious* and *H. sapiens*. Black arrow indicates the conserved 1 nucleotide deletion observed in all Cetacea, previously identified premature stop codons are shown in red.

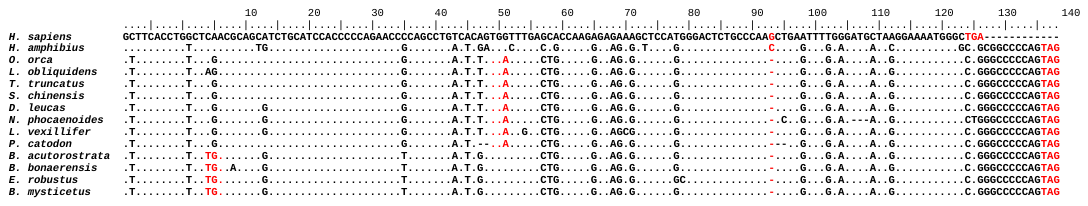
