## Supplementary Material 5 for "Convergent inactivation of the skin-specific C-C motif chemokine ligand 27 in mammalian evolution"

### Slide 1
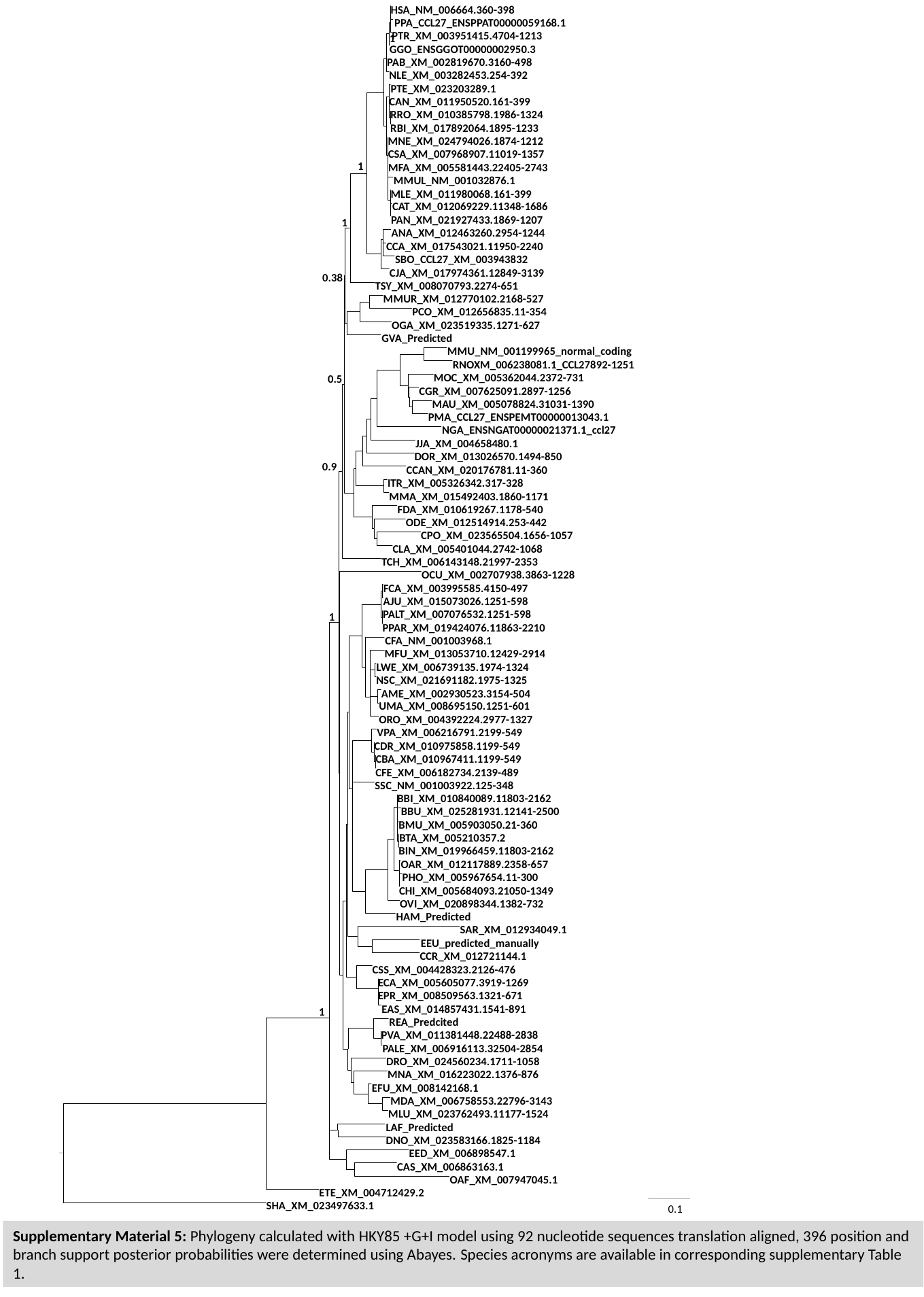

HSA_NM_006664.360-398
PTE_XM_023203289.1
RRO_XM_010385798.1986-1324
CCA_XM_017543021.11950-2240
NGA_ENSNGAT00000021371.1_ccl27
MFU_XM_013053710.12429-2914
LWE_XM_006739135.1974-1324
BMU_XM_005903050.21-360
BIN_XM_019966459.11803-2162
CHI_XM_005684093.21050-1349
OVI_XM_020898344.1382-732
EEU_predicted_manually
MNA_XM_016223022.1376-876
MLU_XM_023762493.11177-1524
CAS_XM_006863163.1
SHA_XM_023497633.1
0.1
PPA_CCL27_ENSPPAT00000059168.1
PTR_XM_003951415.4704-1213
1
GGO_ENSGGOT00000002950.3
PAB_XM_002819670.3160-498
NLE_XM_003282453.254-392
CAN_XM_011950520.161-399
RBI_XM_017892064.1895-1233
MNE_XM_024794026.1874-1212
CSA_XM_007968907.11019-1357
1
MFA_XM_005581443.22405-2743
MMUL_NM_001032876.1
MLE_XM_011980068.161-399
CAT_XM_012069229.11348-1686
PAN_XM_021927433.1869-1207
1
ANA_XM_012463260.2954-1244
SBO_CCL27_XM_003943832
CJA_XM_017974361.12849-3139
0.38
TSY_XM_008070793.2274-651
MMUR_XM_012770102.2168-527
PCO_XM_012656835.11-354
OGA_XM_023519335.1271-627
GVA_Predicted
MMU_NM_001199965_normal_coding
RNOXM_006238081.1_CCL27892-1251
MOC_XM_005362044.2372-731
0.5
CGR_XM_007625091.2897-1256
MAU_XM_005078824.31031-1390
PMA_CCL27_ENSPEMT00000013043.1
JJA_XM_004658480.1
DOR_XM_013026570.1494-850
0.9
CCAN_XM_020176781.11-360
ITR_XM_005326342.317-328
MMA_XM_015492403.1860-1171
FDA_XM_010619267.1178-540
ODE_XM_012514914.253-442
CPO_XM_023565504.1656-1057
CLA_XM_005401044.2742-1068
TCH_XM_006143148.21997-2353
OCU_XM_002707938.3863-1228
FCA_XM_003995585.4150-497
AJU_XM_015073026.1251-598
PALT_XM_007076532.1251-598
1
PPAR_XM_019424076.11863-2210
CFA_NM_001003968.1
NSC_XM_021691182.1975-1325
AME_XM_002930523.3154-504
UMA_XM_008695150.1251-601
ORO_XM_004392224.2977-1327
VPA_XM_006216791.2199-549
CDR_XM_010975858.1199-549
CBA_XM_010967411.1199-549
CFE_XM_006182734.2139-489
SSC_NM_001003922.125-348
BBI_XM_010840089.11803-2162
BBU_XM_025281931.12141-2500
BTA_XM_005210357.2
OAR_XM_012117889.2358-657
PHO_XM_005967654.11-300
HAM_Predicted
SAR_XM_012934049.1
CCR_XM_012721144.1
CSS_XM_004428323.2126-476
ECA_XM_005605077.3919-1269
EPR_XM_008509563.1321-671
EAS_XM_014857431.1541-891
1
REA_Predcited
PVA_XM_011381448.22488-2838
PALE_XM_006916113.32504-2854
DRO_XM_024560234.1711-1058
EFU_XM_008142168.1
MDA_XM_006758553.22796-3143
LAF_Predicted
DNO_XM_023583166.1825-1184
EED_XM_006898547.1
OAF_XM_007947045.1
ETE_XM_004712429.2
Supplementary Material 5: Phylogeny calculated with HKY85 +G+I model using 92 nucleotide sequences translation aligned, 396 position and branch support posterior probabilities were determined using Abayes. Species acronyms are available in corresponding supplementary Table 1.
