## Supplementary Material 1 for "Convergent inactivation of the skin-specific C-C motif chemokine ligand 27 in mammalian evolution"

### Slide 1
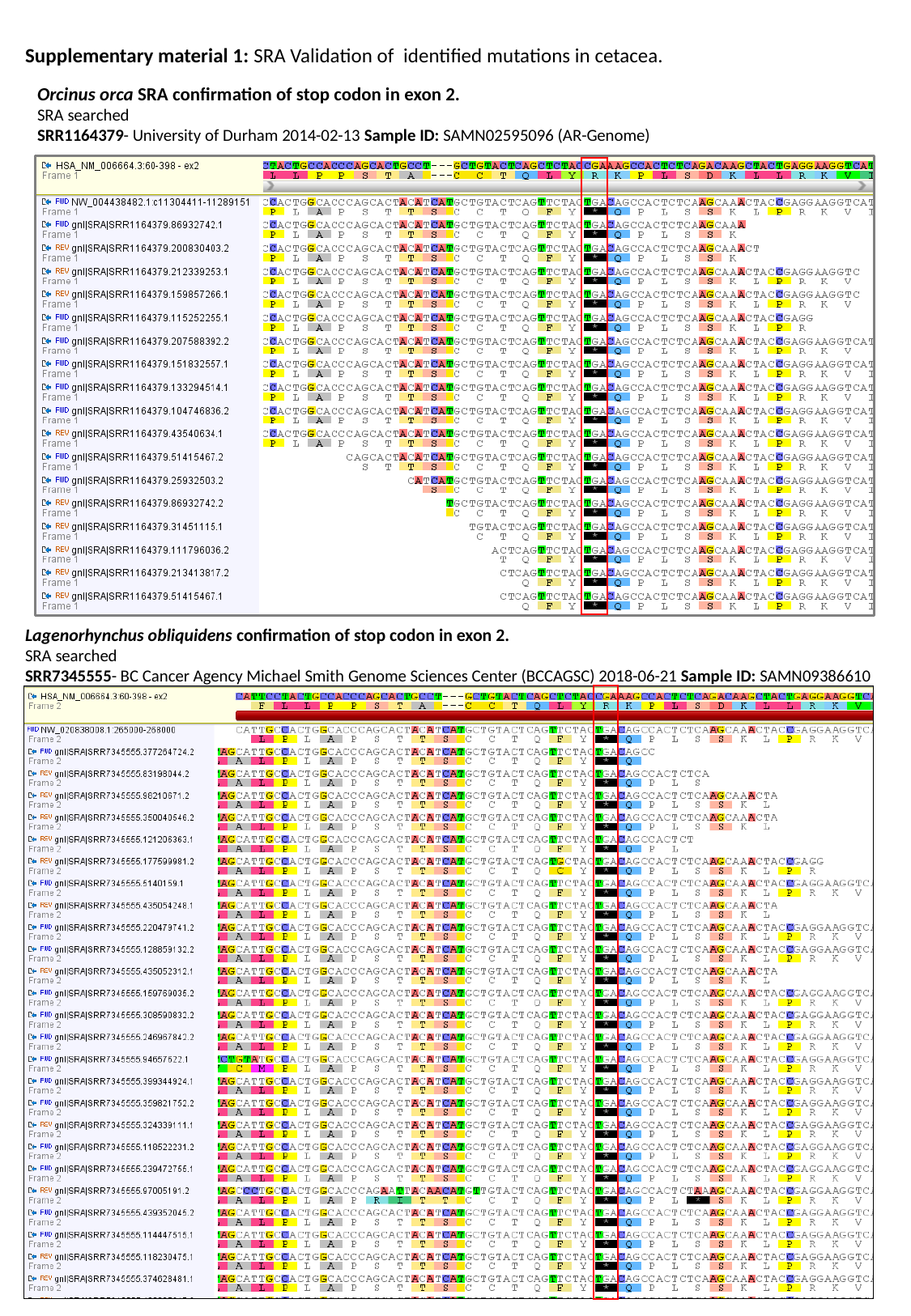

Supplementary material 1: SRA Validation of identified mutations in cetacea.
Orcinus orca SRA confirmation of stop codon in exon 2.
SRA searched
SRR1164379- University of Durham 2014-02-13 Sample ID: SAMN02595096 (AR-Genome)
Lagenorhynchus obliquidens confirmation of stop codon in exon 2.
SRA searched
SRR7345555- BC Cancer Agency Michael Smith Genome Sciences Center (BCCAGSC) 2018-06-21 Sample ID: SAMN09386610

### Slide 2
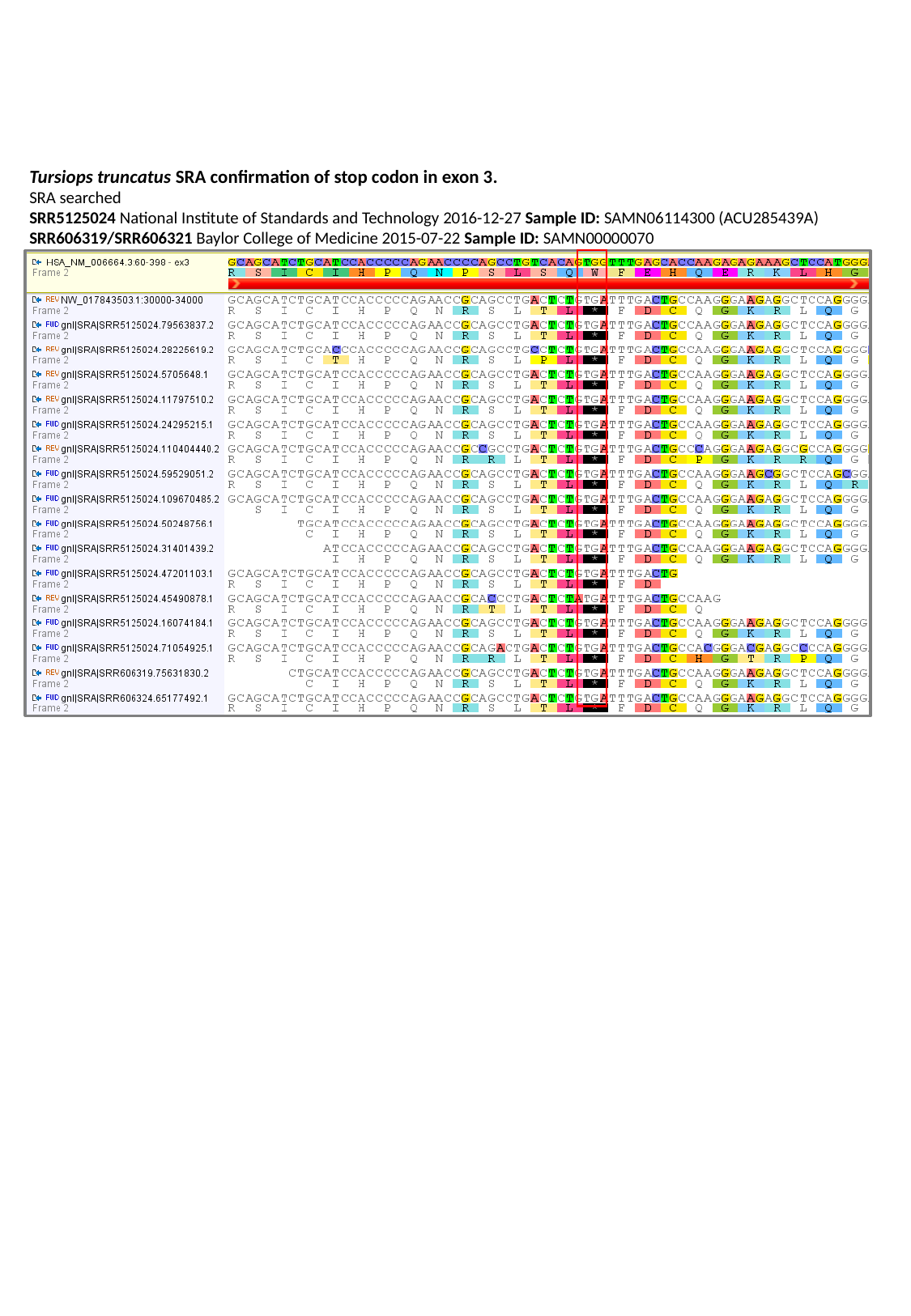

Tursiops truncatus SRA confirmation of stop codon in exon 3.
SRA searched
SRR5125024 National Institute of Standards and Technology 2016-12-27 Sample ID: SAMN06114300 (ACU285439A)
SRR606319/SRR606321 Baylor College of Medicine 2015-07-22 Sample ID: SAMN00000070

### Slide 3
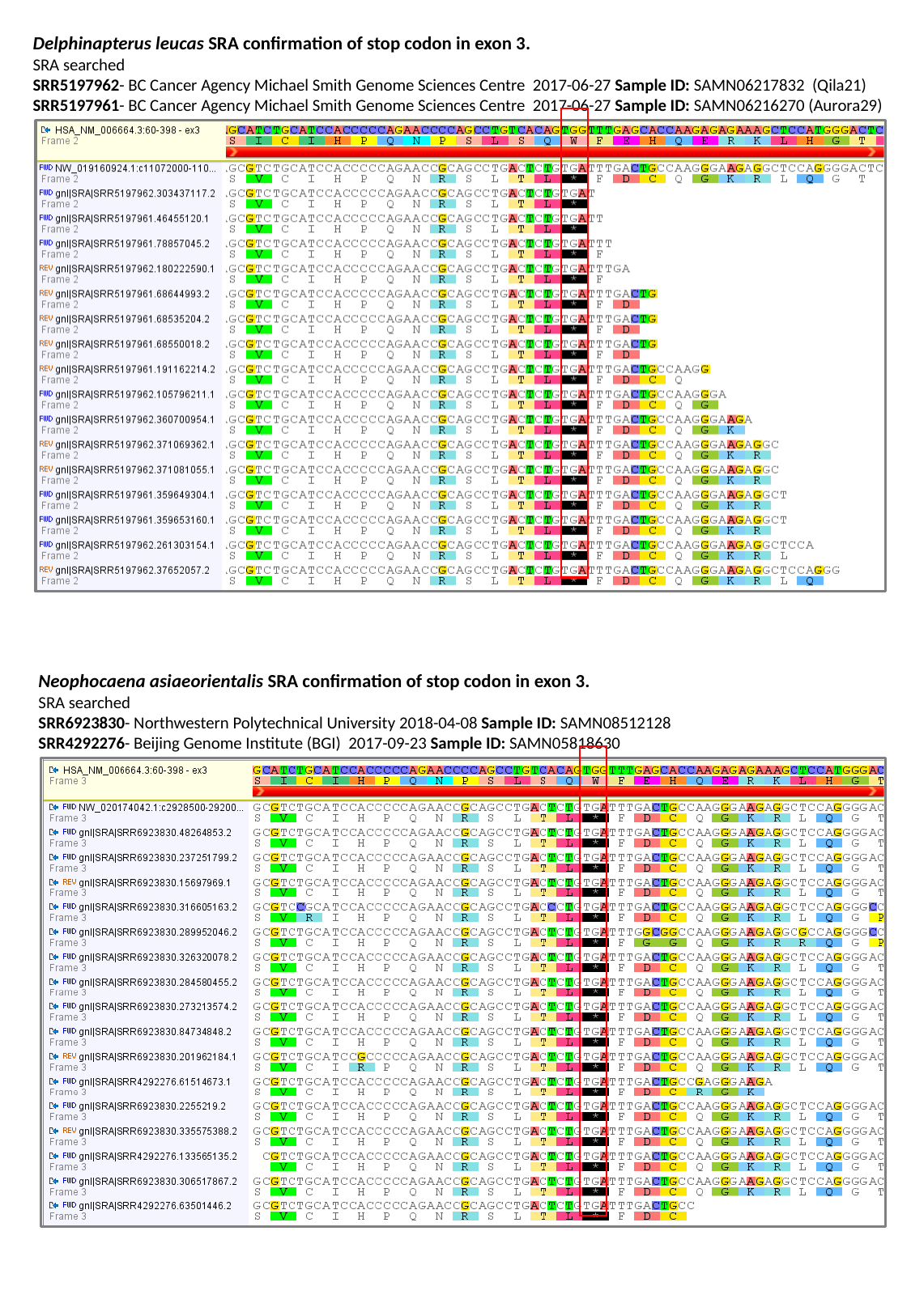

Delphinapterus leucas SRA confirmation of stop codon in exon 3.
SRA searched
SRR5197962- BC Cancer Agency Michael Smith Genome Sciences Centre 2017-06-27 Sample ID: SAMN06217832 (Qila21)
SRR5197961- BC Cancer Agency Michael Smith Genome Sciences Centre 2017-06-27 Sample ID: SAMN06216270 (Aurora29)
Neophocaena asiaeorientalis SRA confirmation of stop codon in exon 3.
SRA searched
SRR6923830- Northwestern Polytechnical University 2018-04-08 Sample ID: SAMN08512128
SRR4292276- Beijing Genome Institute (BGI) 2017-09-23 Sample ID: SAMN05818630

### Slide 4
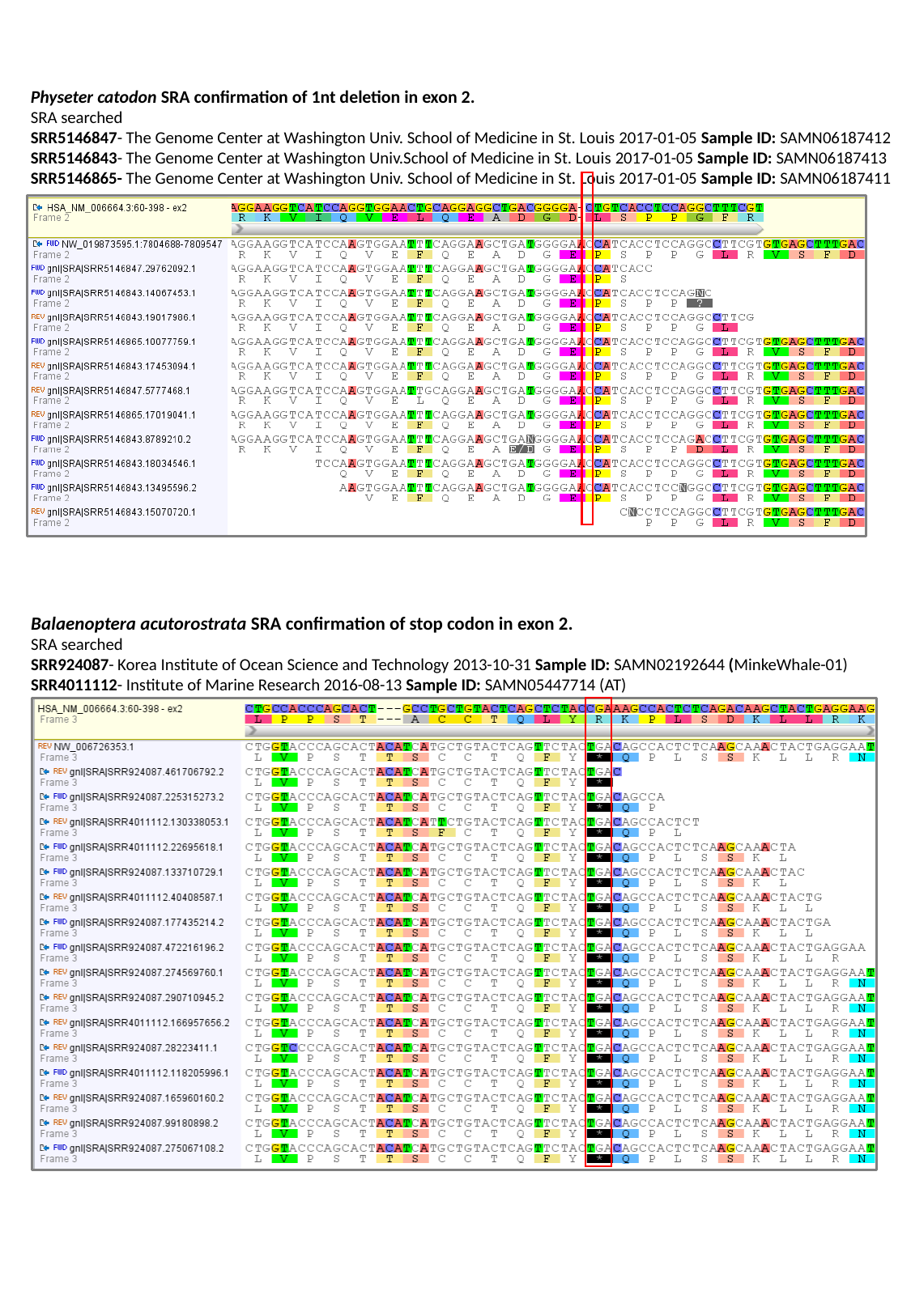

Physeter catodon SRA confirmation of 1nt deletion in exon 2.
SRA searched
SRR5146847- The Genome Center at Washington Univ. School of Medicine in St. Louis 2017-01-05 Sample ID: SAMN06187412
SRR5146843- The Genome Center at Washington Univ.School of Medicine in St. Louis 2017-01-05 Sample ID: SAMN06187413
SRR5146865- The Genome Center at Washington Univ. School of Medicine in St. Louis 2017-01-05 Sample ID: SAMN06187411
Balaenoptera acutorostrata SRA confirmation of stop codon in exon 2.
SRA searched
SRR924087- Korea Institute of Ocean Science and Technology 2013-10-31 Sample ID: SAMN02192644 (MinkeWhale-01)
SRR4011112- Institute of Marine Research 2016-08-13 Sample ID: SAMN05447714 (AT)

### Slide 5
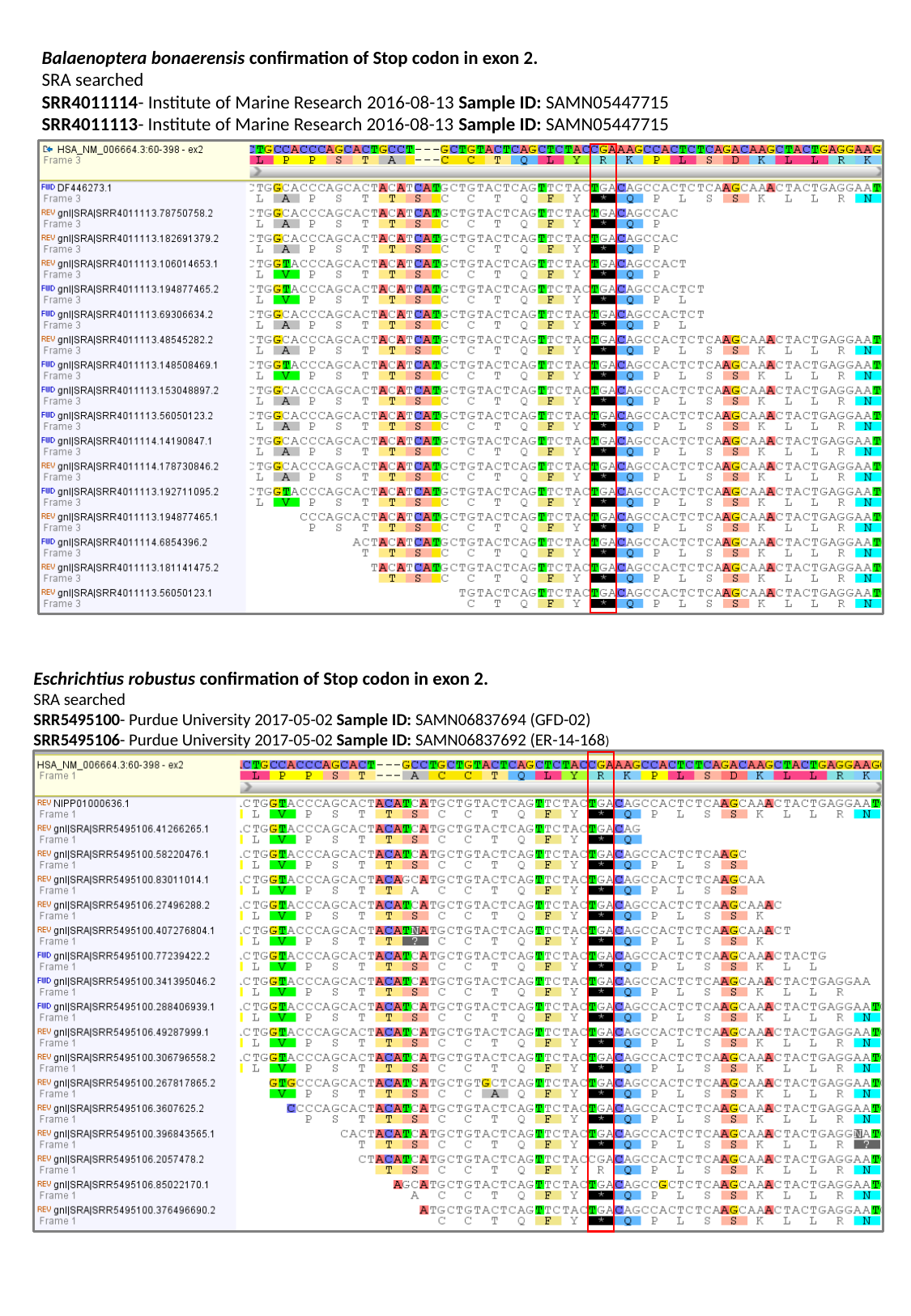

Balaenoptera bonaerensis confirmation of Stop codon in exon 2.
SRA searched
SRR4011114- Institute of Marine Research 2016-08-13 Sample ID: SAMN05447715
SRR4011113- Institute of Marine Research 2016-08-13 Sample ID: SAMN05447715
Eschrichtius robustus confirmation of Stop codon in exon 2.
SRA searched
SRR5495100- Purdue University 2017-05-02 Sample ID: SAMN06837694 (GFD-02)
SRR5495106- Purdue University 2017-05-02 Sample ID: SAMN06837692 (ER-14-168)

### Slide 6
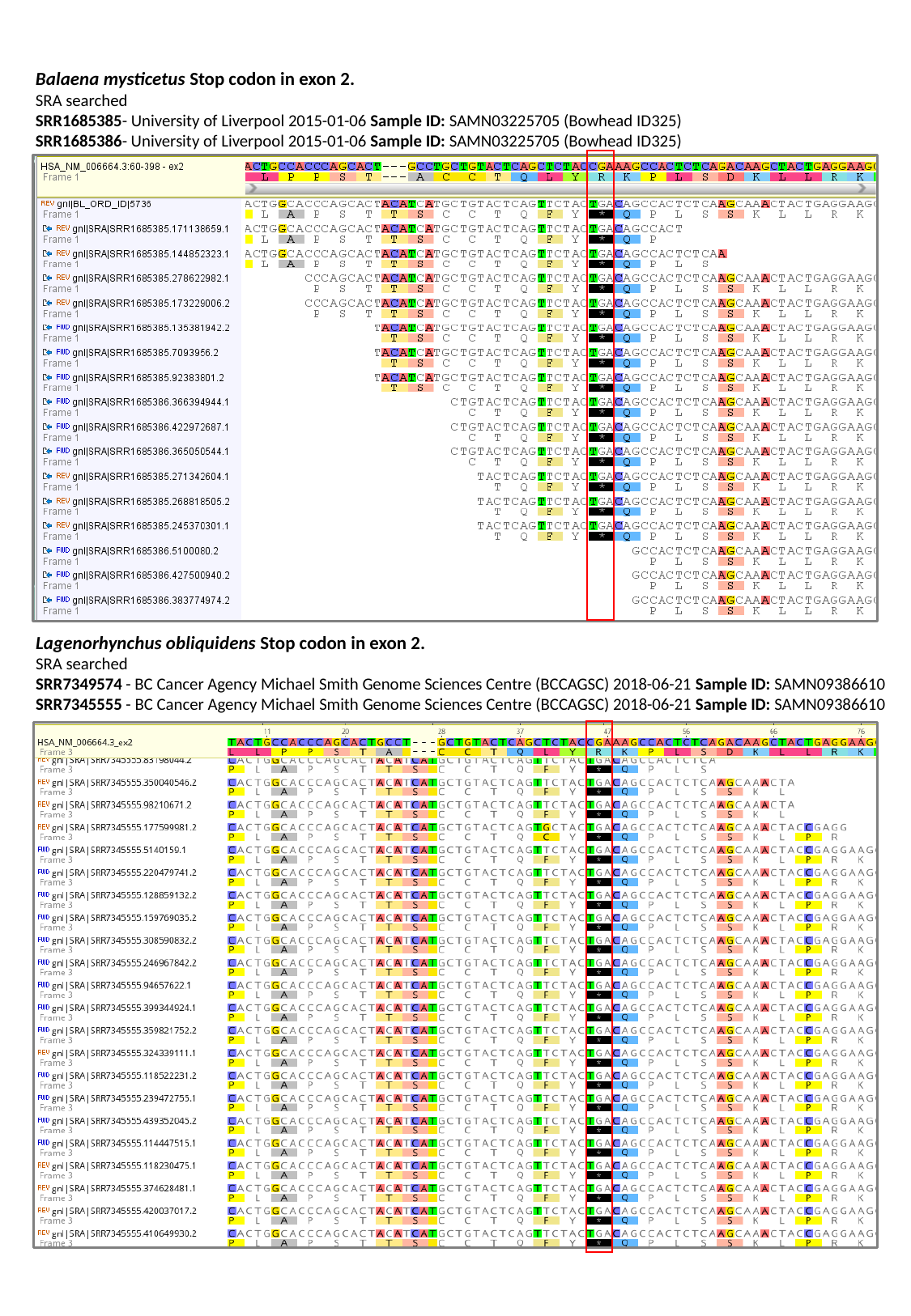

Balaena mysticetus Stop codon in exon 2.
SRA searched
SRR1685385- University of Liverpool 2015-01-06 Sample ID: SAMN03225705 (Bowhead ID325)
SRR1685386- University of Liverpool 2015-01-06 Sample ID: SAMN03225705 (Bowhead ID325)
Lagenorhynchus obliquidens Stop codon in exon 2.
SRA searched
SRR7349574 - BC Cancer Agency Michael Smith Genome Sciences Centre (BCCAGSC) 2018-06-21 Sample ID: SAMN09386610
SRR7345555 - BC Cancer Agency Michael Smith Genome Sciences Centre (BCCAGSC) 2018-06-21 Sample ID: SAMN09386610
