## Supplementary Material 3 for "Convergent inactivation of the skin-specific C-C motif chemokine ligand 27 in mammalian evolution"

### Slide 1
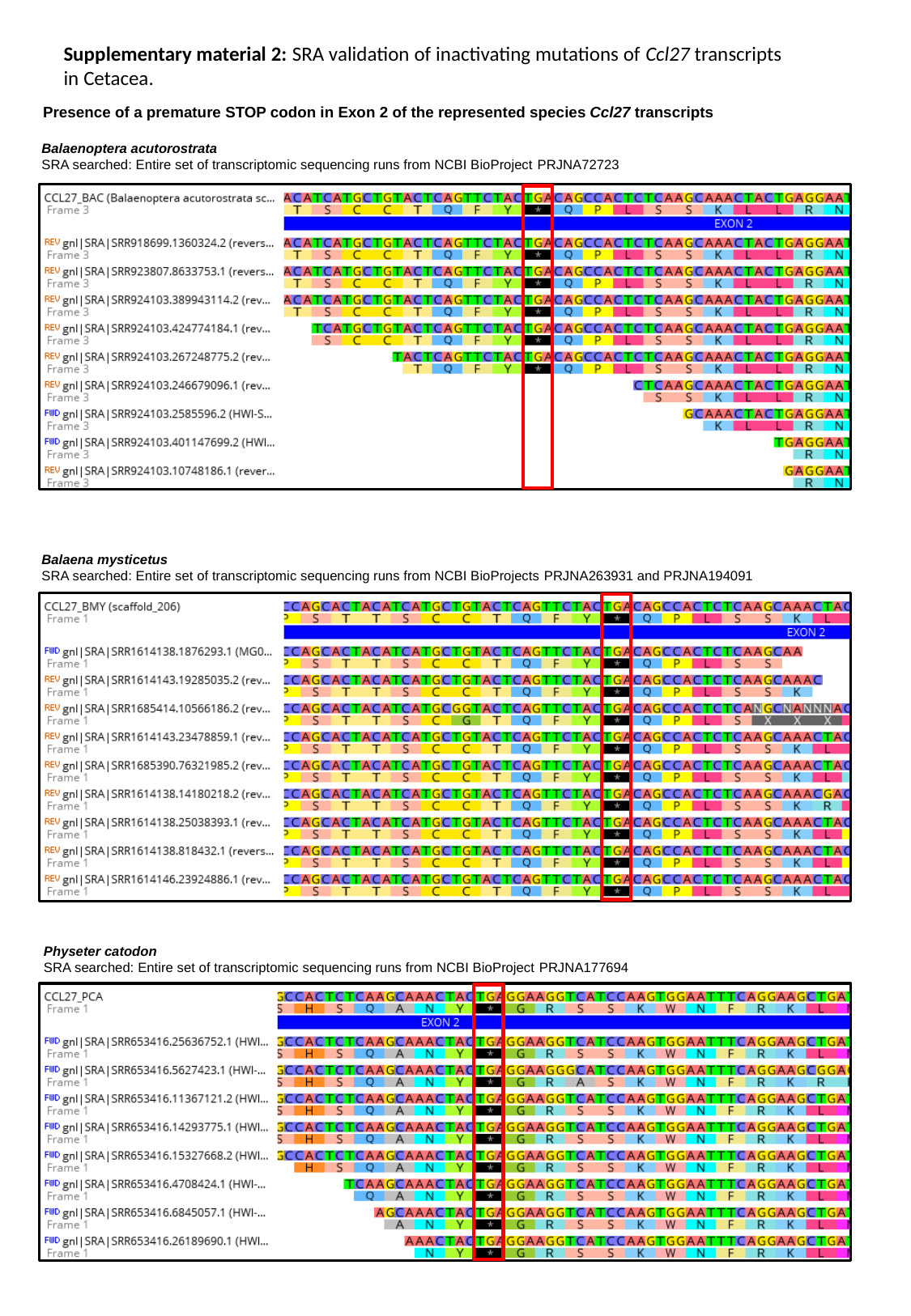

Supplementary material 2: SRA validation of inactivating mutations of Ccl27 transcripts in Cetacea.
Presence of a premature STOP codon in Exon 2 of the represented species Ccl27 transcripts
Balaenoptera acutorostrata
SRA searched: Entire set of transcriptomic sequencing runs from NCBI BioProject PRJNA72723
Balaena mysticetus
SRA searched: Entire set of transcriptomic sequencing runs from NCBI BioProjects PRJNA263931 and PRJNA194091
Physeter catodon
SRA searched: Entire set of transcriptomic sequencing runs from NCBI BioProject PRJNA177694

### Slide 2
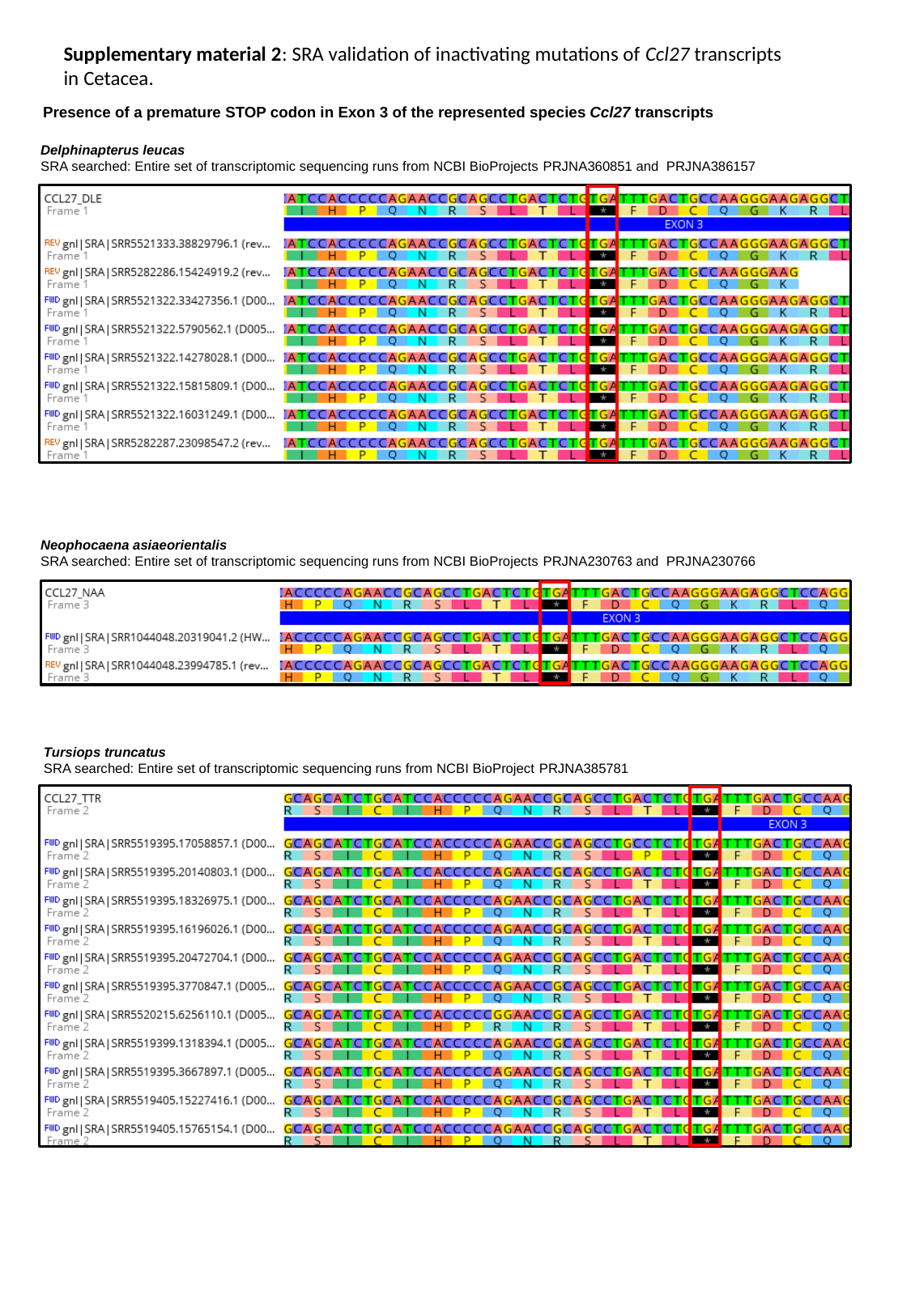

Supplementary material 2: SRA validation of inactivating mutations of Ccl27 transcripts in Cetacea.
Presence of a premature STOP codon in Exon 3 of the represented species Ccl27 transcripts
Delphinapterus leucas
SRA searched: Entire set of transcriptomic sequencing runs from NCBI BioProjects PRJNA360851 and PRJNA386157
Neophocaena asiaeorientalis
SRA searched: Entire set of transcriptomic sequencing runs from NCBI BioProjects PRJNA230763 and PRJNA230766
Tursiops truncatus
SRA searched: Entire set of transcriptomic sequencing runs from NCBI BioProject PRJNA385781
