## Supplementary Table 1 for "Convergent inactivation of the skin-specific C-C motif chemokine ligand 27 in mammalian evolution"

| Supplementary Table 1: Accession numbers of the analysed sequences * tagged low-quality, ^a^ assembled genomes without annotation. | | | | | |
| --- | --- | --- | --- | --- | --- |
| # |  | **Species** | **Order** | **Accession number** | **Exon 3 size** |
| 1 | HSA | *Homo sapiens* | Primate-Hominoidae | NM_006664.3 | 136 |
| 2 | GGO | *Gorilla gorilla gorilla* | Primate-Hominoidae | ENSGGOT00000002950.3 | 136 |
| 3 | PTR | *Pan troglodytes* | Primate-Hominoidae | XM_003951415.4 | 136 |
| 4 | PAB | *Pongo abelii* | Primate-Hominoidae | XM_002819670.3 | 136 |
| 5 | PPA | *Pan paniscus* | Primate-Hominoidae | ENSPPAT00000059168.1 | 136 |
| 6 | NLE | *Nomascus leucogenys* | Primate-Hominoidae | XM_003282453.2 | 136 |
| 7 | MLE | *Mandrillus leucophaeus* | Primate-Cercopithecoidea | XM_011980068.1 | 136 |
| 8 | MNE | *Macaca nemestrina* | Primate-Cercopithecoidea | XM_024794026.1 | 136 |
| 9 | MMUL | *Macaca mulatta* | Primate-Cercopithecoidea | NM_001032876.1 | 136 |
| 10 | MFA | *Macaca fascicularis* | Primate-Cercopithecoidea | XM_005581443.2 | 136 |
| 11 | RRO | *Rhinopithecus roxellana* | Primate-Cercopithecoidea | XM_010385798.1 | 136 |
| 12 | RBI | *Rhinopithecus bieti* | Primate-Cercopithecoidea | XM_017892064.1 | 136 |
| 13 | CAT | *Cercocebus atys* | Primate-Cercopithecoidea | XM_012069229.1 | 136 |
| 14 | CSA | *Chlorocebus sabaeus* | Primate-Cercopithecoidea | XM_007968907.1 | 136 |
| 15 | CAN | *Colobus angolensis palliatus* | Primate-Cercopithecoidea | XM_011950520.1 | 136 |
| 16 | PAN | *Papio anubis* | Primate-Cercopithecoidea | XM_021927433.1 | 136 |
| 17 | PTE | *Piliocolobus tephrosceles* | Primate-Cercopithecoidea | XM_023203289.1 | 136 |
| 18 | ANA | *Aotus nancymaae* | Primate-Platyrrhini | XM_012463260.2 | 88 |
| 19 | SBO | *Saimiri boliviensis** | Primate-Platyrrhini | XM_003943832.2 | 88 |
| 20 | CCA | *Cebus capucinus* | Primate-Platyrrhini | XM_017543021.1 | 88 |
| 21 | CJA | *Callithrix jacchus* | Primate-Platyrrhini | XM_017974361.1 | 88 |
| 22 | OGA | *Otolemur garnettii* | Primate-Strepsirrhini | XM_023519335.1 | 157 |
| 23 | PCO | *Propithecus coquereli* | Primate- Strepsirrhini | XM_012656835.1 | 151 |
| 24 | MMUR | *Microcebus murinus* | Primate- Strepsirrhini | XM_012770102.2 | 157 |
| 25 | TSY | *Tarsius syrichta* | Primate-Haplorrhini | XM_008070793.2 | 157 |
| 26 | TCH | *Tupaia chinensis* | Scandentia | XM_006143148.2 | 157 |
| 27 | GVA | *Galeopterus variegatus** | Dermoptera | XM_008563100.1 | 157 |
| 28 | NGA | *Nannospalax galili* | Rodentia-Myomorpha | ENSNGAT00000021371.1 | 154 |
| 29 | CGR | *Cricetulus griseus* | Rodentia-Myomorpha | XM_007625091.2 | 157 |
| 30 | PMA | *Peromyscus maniculatus bairdii** | Rodentia-Myomorpha | XM_006999053.2*  ENSPEMT00000013043.1 | 157 |
| 31 | MOC | *Microtus ochrogaster* | Rodentia-Myomorpha | XM_005362044.2 | 157 |
| 32 | MMU | *Mus musculus* | Rodentia-Myomorpha | NM_001199965 | 157 |
| 34 | RNO | *Rattus norvegicus* | Rodentia-Myomorpha | XM_006238081.1 | 157 |
| 35 | JJA | *Jaculus jaculus* | Rodentia-Myomorpha | XM_004658480.1 | 163 |
| 36 | MAU | *Mesocricetus auratus* | Rodentia-Myomorpha | XM_005078824.3 | 157 |
| 37 | ITR | *Ictidomys tridecemlineatus* | Rodentia-Sciuromorpha | XM_005326342.3 | 108 |
| 38 | MMA | *Marmota marmota marmota* | Rodentia-Sciuromorpha | XM_015492403.1 | 108 |
| 39 | DOR | *Dipodomys ordii* | Rodentia-Castorimorpha | XM_013026570.1 | 157 |
| 40 | CCAN | *Castor canadensis* | Rodentia-Castorimorpha | XM_020176781.1 | 157 |
| 41 | CPO | *Cavia porcellus* | Rodentia-Hystricomorpha | XM_023565504.1 | 121 |
| 42 | HGL | *Heterocephalus glaber** | Rodentia-Hystricomorpha | XM_004921052.3 ᴪ | 157 |
| 43 | ODE | *Octodon degus* | Rodentia-Hystricomorpha | XM_012514914.2 | 121 |
| 44 | FDA | *Fukomys damarensis* | Rodentia-Hystricomorpha | XM_010619267.1 | 157 |
| 45 | CLA | *Chinchilla lanigera* | Rodentia-Hystricomorpha | XM_005401044.2 | 121 |
| 46 | OCU | *Oryctolagus cuniculus* | Lagomorpha | XM_002707938.3 | 163 |
| 47 | OPR | *Ochotona princeps* | Lagomorpha | Poor genome coverage |  |
| 48 | SSC | *Sus scrofa* | Cetartiodactyla -Suina | NM_001003922.1 | 121 |
| 49 | VPA | *Vicugna pacos* | Cetartiodactyla-Camelidae | XM_006216791.2 | 148 |
| 50 | CFE | *Camelus ferus* | Cetartiodactyla-Camelidae | XM_006182734.2 | 148 |
| 51 | CBA | *Camelus bactrianus* | Cetartiodactyla-Camelidae | XM_010967411.1 | 148 |
| 52 | CDR | *Camelus dromedarius* | Cetartiodactyla-Camelidae | XM_010975858.1 | 148 |
| 53 | BMU | *Bos mutus* | Cetartiodactyla-Bovinae | XM_005903050.2 | 157 |
| 54 | BBI | *Bison bison bison* | Cetartiodactyla-Bovinae | XM_010840089.1 | 157 |
| 55 | BTA | *Bos taurus* | Cetartiodactyla-Bovinae | XM_005210357.2 | 157 |
| 56 | BBU | *Bubalus bubalis* | Cetartiodactyla-Bovinae | XM_025281931.1 | 157 |
| 57 | BIN | *Bos indicus* | Cetartiodactyla-Bovinae | XM_019966459.1 | 157 |
| 58 | OAR | *Ovis aries* | Cetartiodactyla-Caprinae | XM_012117889.2 | 97 |
| 59 | CHI | *Capra hircus* | Cetartiodactyla-Caprinae | XM_005684093.2 | 97 |
| 60 | OVI | *Odocoileus virginianus texanus* | Cetartiodactyla-Cervidae | XM_020898344.1 | 148 |
| 61 | PHO | *Pantholops hodgsonii* | Cetartiodactyla-Antilopinae | XM_005967654.1 | 97 |
| 62 | OOR | *Orcinus orca* | Cetartiodactyla-Cetacea | Not annotated  Manual prediction ᴪ | 61 |
| 63 | TTR | *Tursiops truncatus** | Cetartiodactyla-Cetacea | XM_004313179.2 ᴪ | 61 |
| 64 | LVE | *Lipotes vexillifer** | Cetartiodactyla-Cetacea | XM_007472117.1 ᴪ | 61 |
| 65 | SCH^a^ | *Sousa chinensis* | Cetartiodactyla-Cetacea | QWLN01034924.1 ᴪ | 61 |
| 66 | LOB | *Lagenorhynchus obliquidens** | Cetartiodactyla-Cetacea | XM_027088946.1 ᴪ | 61 |
| 67 | DLE | *Delphinapterus leucas** | Cetartiodactyla-Cetacea | XM_022551649.1 ᴪ | 61 |
| 68 | NAS | *Neophocaena asiaeorientalis* | Cetartiodactyla-Cetacea | XM_024753768.1 ᴪ | 61 |
| 69 | PCA | *Physeter catodon** | Cetartiodactyla-Cetacea | XM_007100747.2 ᴪ | 61 |
| 70 | BAC | *Balaenoptera acutorostrata scammoni* | Cetartiodactyla-Cetacea | No annotation  Manual prediction ᴪ | 16 |
| 71 | BBO^a^ | *Balaenoptera bonaerensis* | Cetartiodactyla-Cetacea | Scaffold DF446273.1 ᴪ | 16 |
| 72 | ERO^a^ | *Eschrichtius robustus* | Cetartiodactyla-Cetacea | NIPP01000636.1 ᴪ | 16 |
| 73 | BMY^a^ | *Balaena mysticetus* | Cetartiodactyla-Cetacea | Scaffold206 ᴪ | 16 |
| 74 | HAM^a^ | *Hippopotamus amphibius* | Cetartiodactyla-Hippopotamidae | Scaffold1867  Manual prediction coding | 148 |
| 75 | CSI | *Ceratotherium simum* | Perissodactyla-Rhinoceratidae | XM_004428323.2 | 148 |
| 76 | ECA | *Equus caballus* | Perissodactyla-Equidea | XM_005605077.3 | 148 |
| 78 | EPR | *Equus przewalskii* | Perissodactyla-Equidea | XM_008509563.1 | 148 |
| 79 | EAS | *Equus asinus* | Perissodactyla-Equidea | XM_014857431.1 | 148 |
| 80 | NSC | *Neomonachus*  *schauinslandi* | Carnivora-Caniformia | XM_021691182.1 | 148 |
| 81 | ORO | *Odobenus rosmarus divergens* | Carnivora-Caniformia | XM_004392224.2 | 148 |
| 82 | LWE | *Leptonychotes weddellii* | Carnivora-Caniformia | XM_006739135.1 | 148 |
| 83 | AME | *Ailuropoda melanoleuca* | Carnivora-Caniformia | XM_002930523.3 | 148 |
| 84 | UMA | *Ursus maritimus* | Carnivora-Caniformia | XM_008695150.1 | 148 |
| 85 | MFU | *Mustela putorius furo* | Carnivora-Caniformia | XM_013053710.1 | 160 |
| 86 | CFA | *Canis lupus familiaris* | Carnivora-Caniformia | NM_001003968.1 | 157 |
| 87 | PALT | *Panthera tigris altaica* | Carnivora- Feliformia | XM_007076532.1 | 145 |
| 88 | FCA | *Felis catus* | Carnivora- Feliformia | XM_003995585.4 | 145 |
| 89 | AJU | *Acinonyx jubatus* | Carnivora- Feliformia | XM_015073026.1 | 145 |
| 90 | PPAR | *Panthera pardus* | Carnivora- Feliformia | XM_019424076.1 | 145 |
| 91 | PVA | *Pteropus vampyrus* | Chiroptera | XM_011381448.2 | 148 |
| 92 | EFU | *Eptesicus fuscus* | Chiroptera | XM_008142168.1 | 157 |
| 93 | PALE | *Pteropus alecto* | Chiroptera | XM_006916113.3 | 148 |
| 94 | MDA | *Myotis davidii* | Chiroptera | XM_006758553.2 | 148 |
| 95 | MNA | *Miniopterus natalensis* | Chiroptera | XM_016223022.1 | 160 |
| 96 | MLU | *Myotis lucifugus* | Chiroptera | XM_023762493.1 | 148 |
| 97 | RAE | *Rousettus aegyptiacus* | Chiroptera | No annotation  Manual prediction coding | 115 |
| 98 | DRO | *Desmodus rotundus* | Chiroptera | XM_024560234.1 | 145 |
| 99 | RSI | *Rhinolophus sinicus* | Chiroptera | No annotation  Manual prediction ᴪ | 148 |
| 100 | HAR | *Hipposideros armiger** | Chiroptera | XM_019624480.1 ᴪ | 16 |
| 101 | LAF | *Loxodonta africana** | Afrotheria-Proboscidea | Annotation LQ  Manual prediction coding | 106 |
| 102 | TMA | *Trichechus manatus latirostris** | Afrotheria-Sirenia | XM_012554400.2 ᴪ | 61 |
| 103 | ETE | *Echinops telfairi* | Afrotheria-Tenrecidae | XM_004712429.2 | 88 |
| 104 | CAS | *Chrysochloris asiatica** | Afrotheria-Chrysochloridae | XM_006863163.1 | 136 |
| 105 | OAF | *Orycteropus afer* | Afrotheria-Tubulidentata | XM_007947045.1 | 160 |
| 106 | EED | *Elephantulus edwardii* | Afrotheria-Macroscelidea | XM_006898547.1 | 148 |
| 107 | MJA | *Manis javanica** | Pholidota | XM_017657544.1 ᴪ | 72 |
| 108 | MPE^a^ | *Manis pentadactyla* | Pholidota | Manual prediction ᴪ | 73 |
| 109 | CCR | *Condylura cristata* | Soricomorpha | XM_012721144.1 | 106 |
| 110 | SAR | *Sorex araneus* | Eulipotyphla | XM_012934049.1 | 148 |
| 111 | EEU | *Erinaceus europaeus* | Erinaceomorpha | No annotation  Manual prediction coding | 106 |
| 112 | DNO | *Dasypus novemcinctus* | Cingulata | XM_023583166.1 | 157 |
| 113 | SHA | *Sarcophilus harrisii* | Marsupialia | XM_023497633.1 |  |
| 114 | MDO | *Monodelphis domestica* | Marsupialia | Not found |  |
