## Supplementary Table 2 for "Convergent inactivation of the skin-specific C-C motif chemokine ligand 27 in mammalian evolution"

| **Species** | **BioProject** | **BioSample** | **Sequencing Run** | **Tissue** | **MBases** | **MBytes** |
| --- | --- | --- | --- | --- | --- | --- |
| *Balaena mysticetus* | PRJNA263931 | SAMN03108659 | SRR1614138 | Kidney | 5472 | 3646 |
|  |  | SAMN03108660 | SRR1614140 | Kidney | 5928 | 3967 |
|  |  | SAMN03108661 | SRR1614142 | Kidney | 4555 | 3047 |
|  |  | SAMN03108662 | SRR1614143 | Kidney | 4959 | 3312 |
|  |  | SAMN03108663 | SRR1614145 | Liver | 4561 | 3034 |
|  |  | SAMN03108664 | SRR1614146 | Liver | 5494 | 3664 |
|  |  | SAMN03108665 | SRR1614166 | Liver | 5339 | 3570 |
|  | PRJNA194091 | SAMN03225706 | SRR1685388 | Muscle | 15178 | 10427 |
|  |  | SAMN03225707 | SRR1685390 | Kidney | 14069 | 9799 |
|  |  | SAMN03225708 | SRR1685413 | Heart | 3178 | 2171 |
|  |  | SAMN03225709 | SRR1685414 | Cerebellum | 2040 | 1393 |
|  |  | SAMN03225710 | SRR1685415 | Liver | 3175 | 2170 |
|  |  | SAMN03225711 | SRR1685416 | Testis | 2526 | 1716 |
|  |  | SAMN03225712 | SRR1685417 | Retina | 11238 | 7569 |
| *Balaenoptera acutorostrata* | PRJNA72723 | SAMN02194641 | SRR918699 | Brain | 4908 | 3410 |
|  |  | SAMN02194642 | SRR918701 | Heart | 8213 | 5721 |
|  |  | SAMN02194643 | SRR919295 | Kidney | 5491 | 3821 |
|  |  | SAMN02194676 | SRR919296 | Liver | 4562 | 3159 |
|  |  | SAMN02194640 | SRR922125 | Lung | 2875 | 2000 |
|  |  | SAMN02194679 | SRR922171 | Muscle | 5592 | 3917 |
|  |  | SAMN02194680 | SRR922446 | Muscle | 9075 | 6336 |
|  |  | SAMN02194681 | SRR923807 | Muscle | 2959 | 2068 |
| *Delphinapterus leucas* | PRJNA360851 | SAMN06375781 | SRR5282283 | Liver | 4338 | 1606 |
|  |  | SAMN06375782 | SRR5282284 | Brain | 4751 | 1762 |
|  |  | SAMN06375781 | SRR5282285 | Liver | 4326 | 1599 |
|  |  | SAMN06375782 | SRR5282286 | Brain | 4853 | 1804 |
|  |  | SAMN06375782 | SRR5282287 | Brain | 4867 | 1813 |
|  |  | SAMN06373807 | SRR5282288 | Liver | 4206 | 1546 |
|  |  | SAMN06375781 | SRR5282289 | Liver | 4338 | 1599 |
|  |  | SAMN06375782 | SRR5282290 | Brain | 4863 | 1803 |
|  |  | SAMN06373807 | SRR5282291 | Liver | 4311 | 1585 |
|  |  | SAMN06373807 | SRR5282292 | Liver | 4310 | 1589 |
|  |  | SAMN06375781 | SRR5282293 | Liver | 4242 | 1563 |
|  |  | SAMN06374824 | SRR5282294 | Brain | 4441 | 1673 |
|  |  | SAMN06374824 | SRR5282295 | Brain | 4339 | 1635 |
|  |  | SAMN06374824 | SRR5282296 | Brain | 4429 | 1672 |
|  |  | SAMN06374824 | SRR5282297 | Brain | 4452 | 1684 |
|  |  | SAMN06373807 | SRR5282298 | Liver | 4301 | 1583 |
|  | PRJNA386157 | SAMN06925939 | SRR5521314 | Blood | 4295 | 2045 |
|  |  | SAMN06925940 | SRR5521315 | Blood | 4938 | 2338 |
|  |  | SAMN06925938 | SRR5521316 | Blood | 4697 | 2235 |
|  |  | SAMN06925937 | SRR5521317 | Blood | 4573 | 2158 |
|  |  | SAMN06925953 | SRR5521318 | Blood | 5014 | 2365 |
|  |  | SAMN06925952 | SRR5521319 | Blood | 5816 | 2748 |
|  |  | SAMN06925954 | SRR5521320 | Blood | 5243 | 2545 |
|  |  | SAMN06925958 | SRR5521321 | Blood | 5358 | 2589 |
|  |  | SAMN06925957 | SRR5521322 | Blood | 5325 | 2577 |
|  |  | SAMN06925956 | SRR5521323 | Blood | 5278 | 2570 |
|  |  | SAMN06925955 | SRR5521324 | Blood | 5509 | 2683 |
|  |  | SAMN06925959 | SRR5521325 | Blood | 5107 | 2484 |
|  |  | SAMN06925960 | SRR5521326 | Blood | 4614 | 2129 |
|  |  | SAMN06925951 | SRR5521327 | Blood | 3271 | 1517 |
|  |  | SAMN06925950 | SRR5521328 | Blood | 5235 | 2422 |
|  |  | SAMN06925949 | SRR5521329 | Blood | 4305 | 2014 |
|  |  | SAMN06925948 | SRR5521330 | Blood | 4963 | 2307 |
|  |  | SAMN06925947 | SRR5521331 | Blood | 5013 | 2333 |
|  |  | SAMN06925946 | SRR5521332 | Blood | 5351 | 2625 |
|  |  | SAMN06925945 | SRR5521333 | Blood | 6845 | 3362 |
|  |  | SAMN06925944 | SRR5521334 | Blood | 5784 | 2835 |
|  |  | SAMN06925943 | SRR5521335 | Blood | 4240 | 2083 |
|  |  | SAMN06925942 | SRR5521336 | Blood | 4451 | 2202 |
|  |  | SAMN06925941 | SRR5521337 | Blood | 4914 | 2410 |
|  | PRJNA360851 | SAMN07509519 | SRR5990716 | Multi-tissue | 4320 | 1877 |
|  |  | SAMN07509506 | SRR5990717 | Lung | 4360 | 1805 |
|  |  | SAMN06216270 | SRR5990718 | Blood | 3878 | 1747 |
|  | PRJNA414234 | SAMN07782284 | SRR6181296 | Skin | 907 | 431 |
|  |  | SAMN07782284 | SRR6181297 | Skin | 889 | 424 |
|  |  | SAMN07782284 | SRR6181298 | Skin | 952 | 443 |
|  |  | SAMN07782284 | SRR6181299 | Skin | 935 | 437 |
|  |  | SAMN07782286 | SRR6181300 | Skin | 818 | 391 |
|  |  | SAMN07782286 | SRR6181301 | Skin | 838 | 399 |
|  |  | SAMN07782286 | SRR6181302 | Skin | 859 | 403 |
|  |  | SAMN07782286 | SRR6181303 | Skin | 881 | 411 |
| *Neophocaena asiaeorientalis* | PRJNA230763 | SAMN02389530 | SRR1044048 | Kidney | 4670 | 2956 |
|  | PRJNA230766 | SAMN02389531 | SRR1044071 | Kidney | 4711 | 2973 |
| *Physeter catodon* | PRJNA177694 | SAMN01906660 | SRR653403 | Not specified | 209 | 127 |
|  |  | SAMN01906658 | SRR653404 |  | 218 | 133 |
|  |  | SAMN01906656 | SRR653405 |  | 241 | 146 |
|  |  | SAMN01906659 | SRR653406 |  | 219 | 133 |
|  |  | SAMN01906657 | SRR653415 |  | 5581 | 4089 |
|  |  | SAMN01906660 | SRR653416 |  | 6131 | 4194 |
|  |  | SAMN01906656 | SRR653419 |  | 6951 | 4751 |
|  |  | SAMN01906659 | SRR653420 |  | 6353 | 4354 |
|  |  | SAMN01906658 | SRR653428 |  | 6338 | 4351 |
| *Tursiops truncatus* | PRJNA385781 | SAMN06909707 | SRR5519395 | Skin | 3491 | 1539 |
|  |  | SAMN06909708 | SRR5519396 | Skin | 3592 | 1517 |
|  |  | SAMN06909717 | SRR5519397 | Skin | 3001 | 1242 |
|  |  | SAMN06909718 | SRR5519398 | Skin | 2765 | 1128 |
|  |  | SAMN06909719 | SRR5519399 | Skin | 3236 | 1313 |
|  |  | SAMN06909720 | SRR5519400 | Skin | 2952 | 1203 |
|  |  | SAMN06909709 | SRR5519401 | Skin | 4436 | 1895 |
|  |  | SAMN06909710 | SRR5519402 | Skin | 3574 | 1530 |
|  |  | SAMN06909711 | SRR5519403 | Skin | 3349 | 1431 |
|  |  | SAMN06909712 | SRR5519404 | Skin | 3297 | 1405 |
|  |  | SAMN06909713 | SRR5519405 | Skin | 2763 | 1172 |
|  |  | SAMN06909714 | SRR5519406 | Skin | 3573 | 1519 |
|  |  | SAMN06909715 | SRR5519407 | Skin | 3202 | 1356 |
|  |  | SAMN06909716 | SRR5519408 | Skin | 3642 | 1488 |
|  |  | SAMN06920712 | SRR5520198 | Skin | 2924 | 1192 |
|  |  | SAMN06920713 | SRR5520199 | Skin | 2790 | 1140 |
|  |  | SAMN06920722 | SRR5520200 | Skin | 3375 | 1408 |
|  |  | SAMN06920723 | SRR5520201 | Blood | 5991 | 2947 |
|  |  | SAMN06920724 | SRR5520202 | Blood | 6650 | 3263 |
|  |  | SAMN06920725 | SRR5520203 | Blood | 6840 | 3365 |
|  |  | SAMN06920726 | SRR5520204 | Blood | 6427 | 3154 |
|  |  | SAMN06920727 | SRR5520205 | Blood | 7139 | 3491 |
|  |  | SAMN06920728 | SRR5520206 | Blood | 6839 | 3397 |
|  |  | SAMN06920729 | SRR5520207 | Blood | 6231 | 3074 |
|  |  | SAMN06920730 | SRR5520208 | Blood | 6492 | 3194 |
|  |  | SAMN06920731 | SRR5520209 | Blood | 6556 | 3231 |
|  |  | SAMN06920714 | SRR5520210 | Skin | 3208 | 1303 |
|  |  | SAMN06920732 | SRR5520211 | Blood | 6000 | 2938 |
|  |  | SAMN06920715 | SRR5520212 | Skin | 3255 | 1360 |
|  |  | SAMN06920716 | SRR5520213 | Skin | 3462 | 1461 |
|  |  | SAMN06920717 | SRR5520214 | Skin | 3158 | 1325 |
|  |  | SAMN06920718 | SRR5520215 | Skin | 3602 | 1507 |
|  |  | SAMN06920719 | SRR5520216 | Skin | 3533 | 1510 |
|  |  | SAMN06920720 | SRR5520217 | Skin | 2991 | 1260 |
|  |  | SAMN06920721 | SRR5520218 | Skin | 3383 | 1423 |

**Supplementary Table 2** – In-depth description of the available transcriptomic NCBI sequence read archive (SRA) projects, scrutinized in the transcriptomic analysis of the 6 represented cetaceans.
